# Characterization of the minimal residual disease state reveals distinct evolutionary trajectories of human glioblastoma

**DOI:** 10.1101/2022.01.28.478232

**Authors:** Maleeha A. Qazi, Sabra K. Salim, Kevin R. Brown, Nicholas Mickolajewicz, Neil Savage, Hong Han, Minomi K. Subapanditha, David Bakhshinyan, Allison Nixon, Parvez Vora, Kimberly Desmond, Chirayu Chokshi, Mohini Singh, Amanda Khoo, Andrew Macklin, Shahbaz Khan, Nazanin Tatari, Neil Winegarden, Laura Richards, Trevor Pugh, Nicholas Bock, Alireza Mansouri, Chitra Venugopal, Thomas Kislinger, Sidhartha Goyal, Jason Moffat, Sheila K. Singh

## Abstract

Recurrence of solid tumors renders patients vulnerable to a distinctly advanced, highly treatment-refractory disease state that has an increased mutational burden and novel oncogenic drivers not detected at initial diagnosis. Improving outcomes for recurrent cancers requires a better understanding of cancer cell populations that expand from the post-therapy, minimal residual disease (MRD) state. We profiled barcoded tumor stem cell populations through therapy at tumor initiation/engraftment, MRD and recurrence in our therapy-adapted, patient-derived xenograft models of glioblastoma (GBM). Tumors showed distinct patterns of recurrence in which clonal populations exhibited either an *a priori*, pre-existing fitness advantage, or *a priori* equipotency fitness acquired through therapy. Characterization of the MRD state by single-cell and bulk RNA sequencing revealed a tumor-intrinsic immunomodulatory signature with strong prognostic significance at the transcriptomic level and in proteomic analysis of cerebrospinal fluid (CSF) collected from GBM patients at all stages of disease. Our results provide insight into the innate and therapy-driven dynamics of human GBM, and the prognostic value of interrogating the MRD state in solid cancers.

## Introduction

Intratumoral heterogeneity (ITH) as a dynamic phenomenon represents a key determinant of therapy failure across various solid cancers. The continual evolution of genetic, cellular and functional cancer heterogeneity over time and throughout therapy has confounded clinical management of disease^1-3^. The persistence of treatment-resistant subpopulations of cancer stem cells after initial therapy^4^ and their subsequent expansion may, in fact, drive a highly aggressive, heterogeneous and biologically distinct recurrent tumor. An archetype of ITH driving profound treatment resistance can be found in glioblastoma (GBM), the most common primary malignant brain tumor in adults^5-7^. Despite aggressive multimodal treatment with surgical resection, chemotherapy with temozolomide (TMZ) and radiotherapy, no further standard-of-care (SoC) treatment options exist upon recurrence, and only 10% of all GBM patients participate in clinical trials^8^. Recent genomic studies have shown that therapy can act as a selection pressure or bottleneck for tumor evolution from minority cell populations present at the time of initial tumor diagnosis^9-11^. Although current preclinical models fail to predict the recurrent tumor landscape, this roadblock may be circumvented through development of a biological understanding of the small subpopulation of rare, treatment-resistant cells that drive the emerging recurrence.

A critical limitation to our biological understanding of GBM recurrence lies in the scarcity of tissue specimens to profile and study, as only 25% of progressive or recurrent patients undergo repeat surgical resection^12^. We developed methods to generate unique patient-derived xenograft (PDX) models of human GBM recurrence, by adapting SoC chemoradiotherapy in immunocompromised mice intracranially engrafted with patient-derived primary glioma stem cells (GSCs). This method allows for temporal profiling of the clonal and transcriptomic composition of cancer through different stages toward the acquisition of treatment resistance, and especially led to the identification of disease stage characterized as the lowest possible histological and radiological disease burden, which we termed “minimal residual disease” (MRD). This model thus affords an unprecedented window into a highly informative stage, which is currently not possible to interrogate in brain tumors. As MRD has offered invaluable clinical insight into the management of both recurrent liquid cancers^13^ and solid cancers^14-15^, it could offer great therapeutic insights in anticipating the composition of GBM recurrence.

In this work, we combine our unique models of human GBM recurrence with cell tracking using DNA barcodes to examine clonal GSC populations through SoC chemo- and radiotherapy to determine whether recurrence arises stochastically or from pre-existing resistors. Thus, we compare the clonal dynamics of GBM recurrence across multiple patient tumors models of recurrence. Through cell tracking using DNA barcodes and bulk and single-cell RNA sequencing (RNAseq), we characterize the MRD state in human GBM and uncover its potential predictive value in anticipating cancer recurrence.

### Development of xenograft model of GBM recurrence

To characterize tumor evolution through therapy, we developed a therapy-adapted, patient-derived xenograft model to track tumor cells from five primary, treatment-naïve human GBMs (**Supp Table 1-2**). We monitored tumor growth in mice using magnetic resonance imaging (MRI) and began *in vivo* chemoradiotherapy upon visible engraftment (ENG), which led to an overall survival advantage (**Supp Table 3**). Approximately two weeks post-therapy, we observed improved clinical symptoms in treated mice, coupled with minimal radiographic evidence of disease, as is frequently observed in patients. We termed this state minimal residual disease (MRD), a clinically- and biologically-relevant stage of disease comprising the treatment-resistant pool of cells that comprises the imminent recurrence leading to disease end-point (REC) (**Fig 1a**). Phenotypic assessment of tumor cells collected at REC showed increased stem cell frequency and sphere forming capacity after treatment (**Supp Fig 1a-b**), which we have previously shown is associated with poor clinical outcome and prognosis^16-17^. While phenotypic assessments provide insight into the changes of bulk tumor populations through therapy, we sought to assess the effect of therapy on individual clonal populations over the course of disease.

**Figure 1.**
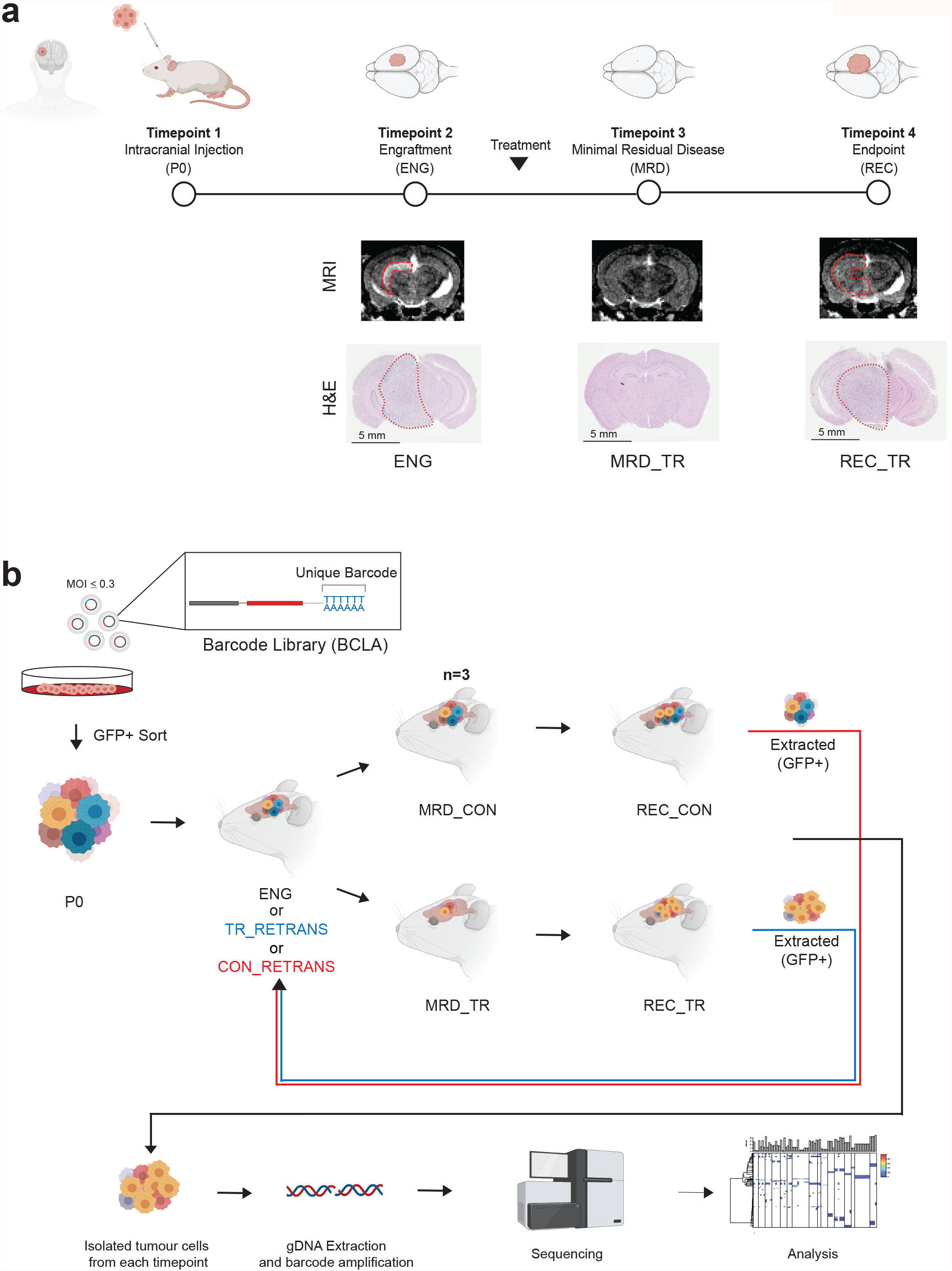
Human-mouse glioblastoma xenograft modeling through chemoradiotherapy regimens. **a**. Experimental timepoints tracking glioblastoma volume through magnetic resonance imaging. **b**. Schematic of tracking glioblastoma evolution through barcoding and serial retransplantation. Briefly, five patient tumors were cultured and infected with the BCLA barcode library. Barcoded cells were sorted by flow cytometry for GFP+ and expanded prior to engraftment (P0). Cells were engrafted into NSG mice, which were culled at the MRD timepoint, or at recurrence after undergoing chemoradiotherapy or placebo treatment (N=3, each arm). Recurrent control and treated tumors of BT799 and BT428, respectively, were extracted and transplanted into a new cohort of mice upon which they completed a second round of *in vivo* expansion with or without treatment for barcode analysis.

### GBM displays innate and therapy-driven clonal dynamics through disease progression

To investigate the clonal changes of GBM through therapy, we adapted a DNA barcoding strategy to track individual tumor cells at multiple timepoints during the course of disease progression and treatment (**Fig 1b**). Tumor cells were transduced with a high-complexity lentiviral barcode library at low multiplicity of infection (MOI), sorted by GFP expression (**Supp Fig 1c**), and expanded prior to intracranial transplantation in immunodeficient mice to ensure accurate tracing of clonal cellular populations. All tumor samples showed high barcode diversity at P0, ensuring many clonal GBM populations were followed. As expected, clonal diversity decreased significantly from P0 to ENG (**Fig 2a-b, Sup Fig 2a-b, left panels**). Engraftment efficiency, as measured by the change in barcode abundance, ranged from 7% (BT935) to 20% (MBT06) and closely matched the expected GSC frequency within our GBMs, as assessed by limiting dilution assay^17^ (**Sup Fig 1a-b**). Across all samples, changes in clonal diversity from ENG to REC in untreated mice (REC_CON) remained relatively constant. However, clonal diversity of different samples responded variably to treatment, as exemplified by BT428 and BT799. While both samples showed an overall decrease in diversity, the effect was greater in BT428 when treated with chemoradiotherapy (REC_TR) (**Fig 2a, left panel**). Treatment of BT428 tumors induced a shift towards clonality, where fewer clones comprised the bulk of the REC_TR tumors. A mean of 11 clones comprised 98% of the treated tumors [min=3, max=18], compared to an average of 3089 clones making up 98% of the control tumors [min=1774, max=5202] (**Fig 2a, right panel**). However, despite a slight decrease in barcode diversity in BT799 (**Fig 2b, left panel**), we did not observe a similar concomitant decrease in total barcodes after treatment. BT799 control tumors had an average of 110 clones making up 98% of the tumor [min=46, max=207], while treated tumors had an average of 89 clones [min=3, max=254]. Assessment of clonal frequency revealed only slight differences between control and treated samples at recurrence (**Fig 2b, right panel**). While BT428 tumors had higher diversity and clonal composition in untreated relative to treated samples, the clonal composition of BT799 was similar irrespective of treatment. Closer inspection of the barcode distributions before and after treatment suggests that BT428 has a more heterogeneous makeup, with a larger number of treatment-sensitive clones that are lost due to treatment, and a smaller fraction of treatment-refractory barcoded cells. In contrast, there is little change observed in the distributions of BT799 untreated and treated populations, supporting a more homogenous treatment-refractory population.

**Figure 2.**
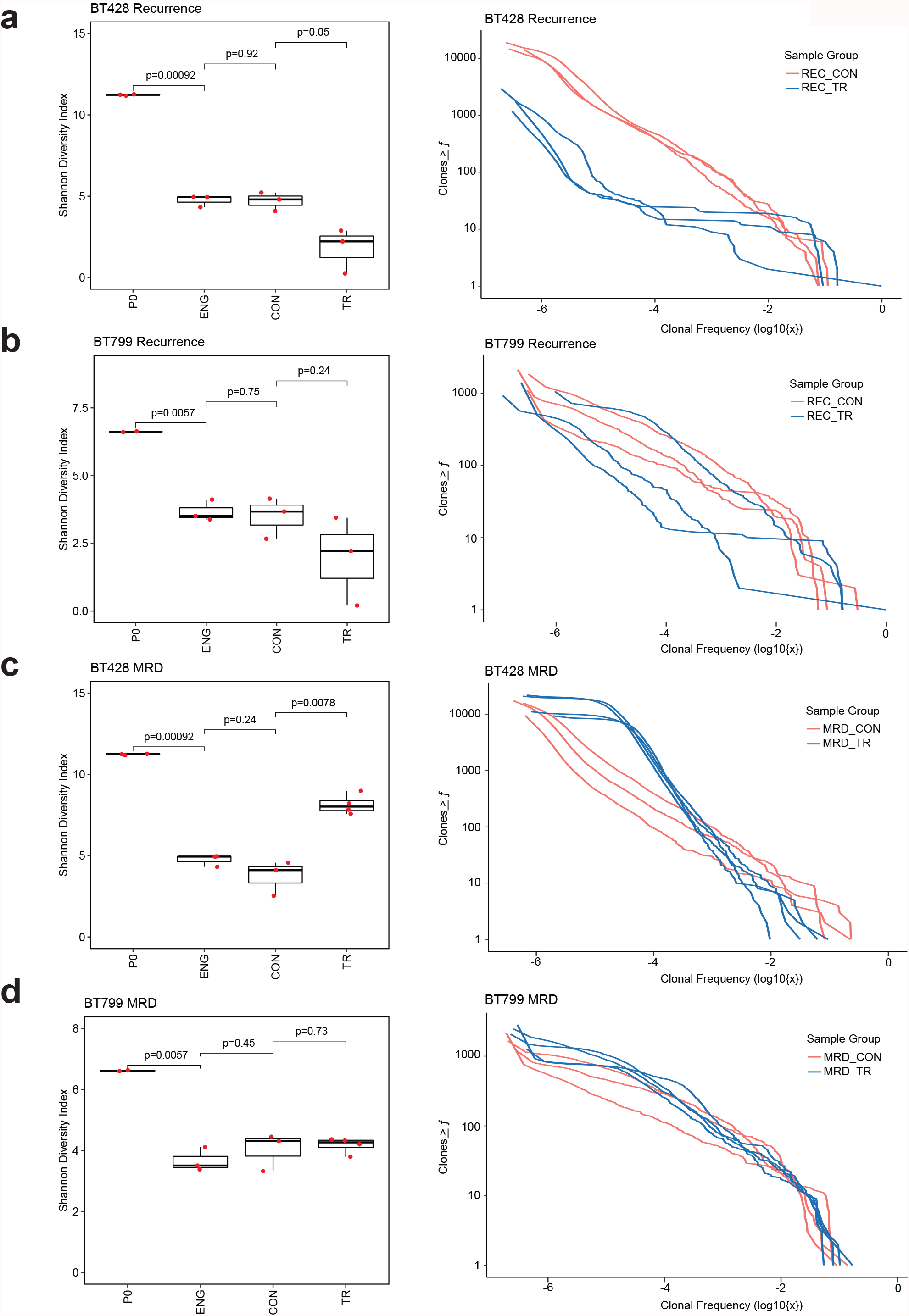
Tracking subclonal frequency of GBM through chemoradiotherapy selective pressures. **a-b**. For BT428 (a) and BT799 (b), shannon Index (left) displays relative diversity of barcodes through timepoints of *in vivo* modeling and clonal frequency (right) at the time of recurrence in response to chemoradiotherapy (blue) and time-matched controls (red). Each curve represents an independent experimental replicate (N=3 for all except N=2 for BT799 P0). P-values determined using Welch Two Sample t-test. **c-d**. For BT428 (c) and BT799 (d), shannon Index (left) displays relative diversity of barcodes through timepoints of *in vivo* modeling and clonal frequency (right) at the time of minimal residual disease (MRD) in response to chemoradiotherapy (blue) and time-matched controls (red). Each curve represents an independent experimental replicate (N=3 for all except N=2 for BT799 P0). P-values determined using Welch Two Sample t-test.

We next sought to define whether clonal composition at MRD could predict patterns of recurrence. Assessment of clonal diversity in BT428 showed an opposing trend at MRD in which treated samples showed significantly increased clonal diversity in comparison to untreated samples (MRD_TR vs MRD_CON, p=0.0078, Welch Two Sample t-test, N = 3 biologically independent replicates) (**Fig 2c, left panel**). Thus, BT428 at MRD showed high clonal diversity even after treatment, while untreated samples showed slight enrichment of low abundance barcodes. Barcode diversity in BT799 showed little change with treatment at MRD (MRD_CON vs MRD_TR, p=0.73, Welch Two-sample t-test) (**Fig 2d, left panel**). Untreated and treated samples showed comparable diversity to that observed at engraftment, suggesting similar clonal kinetics irrespective of treatment. A similar trend was observed with re-transplantation of CON BT799 at endpoint, and subsequent treatment in which the same clonal kinetics were reproducibly observed (**Supp Fig 2d**). Unlike BT428, BT799 showed a shift to clonality at MRD in both treated and time-matched untreated samples, which was similar to what was observed at REC (**Fig 2d, right panel**). Thus, while BT428 and BT799 at REC showed reduced diversity and highly clonal populations with treatment, BT428 appeared to have acquired this diversity between MRD and REC. This suggested that clones that would eventually dominate acquired treatment-resistance, and then rapidly expanded to recurrence. This observation was bolstered by re-transplantation of BT428_REC_TR, which upon additional chemoradiotherapy, revealed little change in clonal composition, suggesting an acquired-treatment resistance and tumorigenicity (**Supp Fig 2e**). In contrast, BT799 appeared to consistently display clonal dominance that was slightly enriched with treatment and retransplantation (BT799_CON_TR)

The quantitative observations of clonal complexity and expansion from MRD to REC suggested two possible mechanisms of recurrence – one in which a non-reproducible, low abundance clone rapidly expands post-treatment (hereafter referred to as *a priori* equipotency), or one in which a pre-existing clone of high abundance remains abundant through disease progression, survives treatment, and continues to expand thereafter (hereafter referred to as *a priori* fitness). To assess these hypotheses, we qualitatively tracked independent clones at MRD and REC. Bubble plots, in which each bubble represents a unique barcode, showed little similarity between replicates across all timepoints in BT428 (**Fig 3a**). In the absence of treatment, few distinct barcodes were reproducibly enriched from ENG to MRD or REC. Similarly, treatment did not reproducibly select distinct barcodes, and barcodes of high abundance at MRD did not remain so at REC (**Fig 3c**). Additionally, abundant clones at ENG did not remain abundant, suggesting against the possibility of clones with pre-existing tumorigenic and treatment-resistant potential. Thus, when combined with the quantitative assessments, these data suggest that clones in BT428 were stochastically enriched from MRD to REC, and that their abundance at MRD did not predict their subsequent expansion.

**Figure 3.**
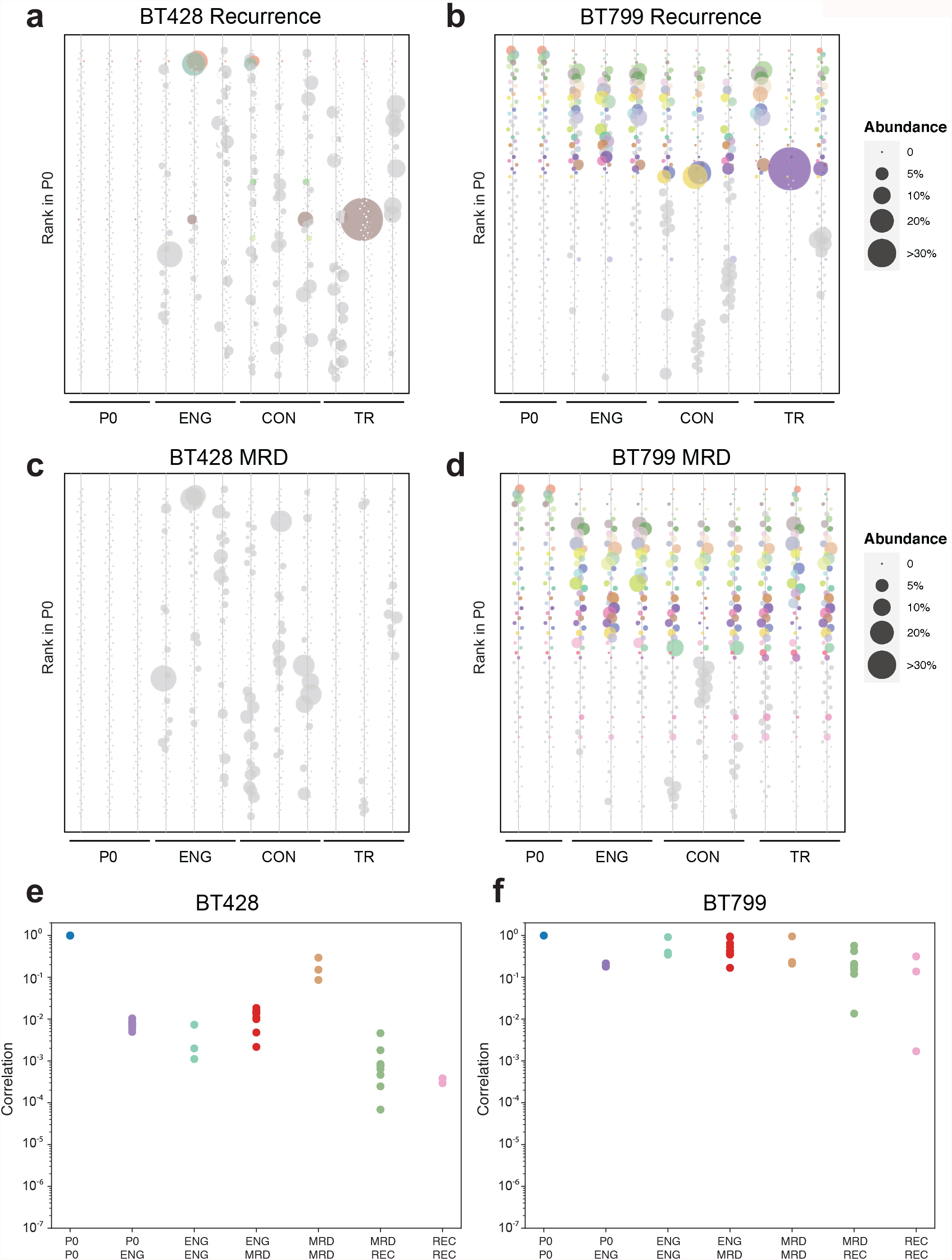
Subclonal frequency of GBMs change over time in response to chemoradiotherapy. **a-b**. Bubble plot displaying frequency of barcodes in (a) BT428 and (b) BT799 at the time of recurrence. Barcodes that enrich in more than one sample (>1%) are colored while barcodes enriched in a single sample are represented in grey. Each vertical line represents an independent experimental replicate for the timepoint (N=3 for all except N=2 for BT799 P0). **c-d**. Bubble plot displaying frequency of barcodes in (c) BT428 and (d) BT799 at the minimal residual disease (MRD) timepoint. Barcodes that enrich in more than one sample (>1%) are colored while barcodes enriched in a single sample are represented in grey. Each vertical line represents an independent experimental replicate for the timepoint (N=3 for all except N=2 for BT799 P0). **e-f**. Barcode correlation of timepoints throughout *in vivo* chemoradiotherapy regimens in (e) BT428 and (f) BT799.

In contrast, BT799 reproducibly showed selection of a few subsets of clones at MRD and REC (**Fig 3b, d**). Clones that were enriched prior to injection (P0) remained abundant from ENG to MRD to REC. While highly abundant clones could have maintained their abundance from *in vitro* expansion simply due to tumor growth *in vivo*, their continued dominance through treatment suggests a pre-existing potential to survive not only *in vitro* conditions (as demonstrated by abundance at P0) but also *in vivo* chemoradiotherapy. Unlike BT428, the clonal composition of BT799 at MRD in both treated and untreated samples later predicted clonal composition at REC. While quantitative assessment of BT799 at REC revealed a slight shift towards clonality with treatment, examination of barcode identity suggests that this was likely due to elimination of lower-abundance, treatment-sensitive clones (**Fig 3b, d**).

To define patterns of recurrence in BT428 and BT799 after treatment, we analyzed barcode identity temporally throughout disease progression. By combining identity and abundance, we assessed correlation of each replicate tumor sample against one another collected at the sample time point and with samples from other timepoints (ex: P0 vs ENG, ENG vs ENG). BT428 and BT799 showed highly correlated starting populations between replicate tumor samples, indicating that downstream changes were due to differential clonal behaviour *in vivo* (P0 vs P0) (**Fig 3e-f**). At ENG, both BT428 and BT799 showed a drop in similarity from the initial starting population, likely due to elimination of clones without tumorigenic potential. However, correlations of samples from ENG onwards confirmed earlier hypotheses of clonal composition through time in BT428 and BT799. BT799 showed high correlation of replicates between each other at ENG and between ENG and MRD (**Fig 3f**). This phenomenon of high intra- and intertemporal correlation was also observed between MRD and REC (ρ ≤ 10^−2.5^), respectively. In contrast, there was a lower correlation between BT428 samples at ENG, and between ENG and MRD, suggesting inconsistent clonal expansion prior to treatment (**Fig 3e**). Interestingly, BT428 samples at MRD were highly correlated across replicate tumor samples, likely due to reproducible elimination of treatment-sensitive clones. However, weak correlation between samples from MRD and REC and between REC samples suggests differential selection of the remaining treatment-resistant clones. These observations were confirmed in a separate analysis using only highly abundant, resistant clones in BT428 and BT799 between replicates and timepoints, in which dominant resistant clones were highly correlated in BT799 and weakly correlated in BT428 across timepoints (**Supp Fig 3j-k)**. These results suggest that BT799 consists of pre-existing, highly fit and treatment-resistant oligoclonal populations that dominate through disease progression, while BT428 may have clones of equipotent expansion potential that are sporadically selected after treatment and acquire treatment resistance.

These two patterns were observed in our other patient-derived samples, where BT935 and MBT06 displayed similar dynamics to BT799, and BT954 was most similar to BT428 (**Supp Fig 2-3**). Interestingly, initial analysis of BT954 suggested a small subset of clones exhibited increased *a priori* fitness, as they remained highly abundant throughout disease progression. However, closer inspection of the barcode identities revealed inconsistent clonal selection between replicates (Supp Fig 3-1c,d), suggesting that numerous clonal subpopulations may have had elevated pre-existing fitness and were randomly selected at ENG.

Taken together, we identified two patterns of GBM recurrence within a cohort of barcoded patient-derived GBMs subjected to our adapted mouse treatment protocol, one in which MRD can predict clonal selection due to *a priori* fitness, and one in which clones exhibit *a priori* equipotency. While these may be distinct patterns of recurrence, observations in BT954 suggest that *a priori* fitness and *a priori* equipotency may exist on a spectrum. We next sought to define whether profiling the biology of MRD could provide further clinical and therapeutic insights.

### Single-cell sequencing of MRD reveals differential transcriptional signatures and intertumoral heterogeneity

To characterize the MRD stage, we performed single-cell RNA sequencing on *in vivo-* modelled, patient-derived lines MBT06 and BT799. While these samples displayed similar clonal kinetics, MBT06 and BT799 represented tumors derived from patients with the best and worst disease outcomes in our cohort (>43 months and 3-month survival, respectively). Thus, we sought to explore differences with therapeutic and/or prognostic relevance.

We profiled the transcriptome of 2,454 tumor cells using scRNAseq from control or treated MBT06 and BT799 mice (**Fig 4a**; BT799_MRD_CON: 1147 cells, BT799_MRD_TR: 126 cells, MBT06_MRD_CON: 628 cells, MBT06_MRD_TR: 553 cells). Comparison of the scRNAseq data with a more stringent subset obtained by high gene recovery (>3000 genes/cell) and low mitochondrial content (<10%) filters (**Supp Fig 4a**) and also with bulk RNAseq profiles (**Supp Fig 4b**) confirmed the high quality of data used for subsequent analyses.

**Figure 4.**
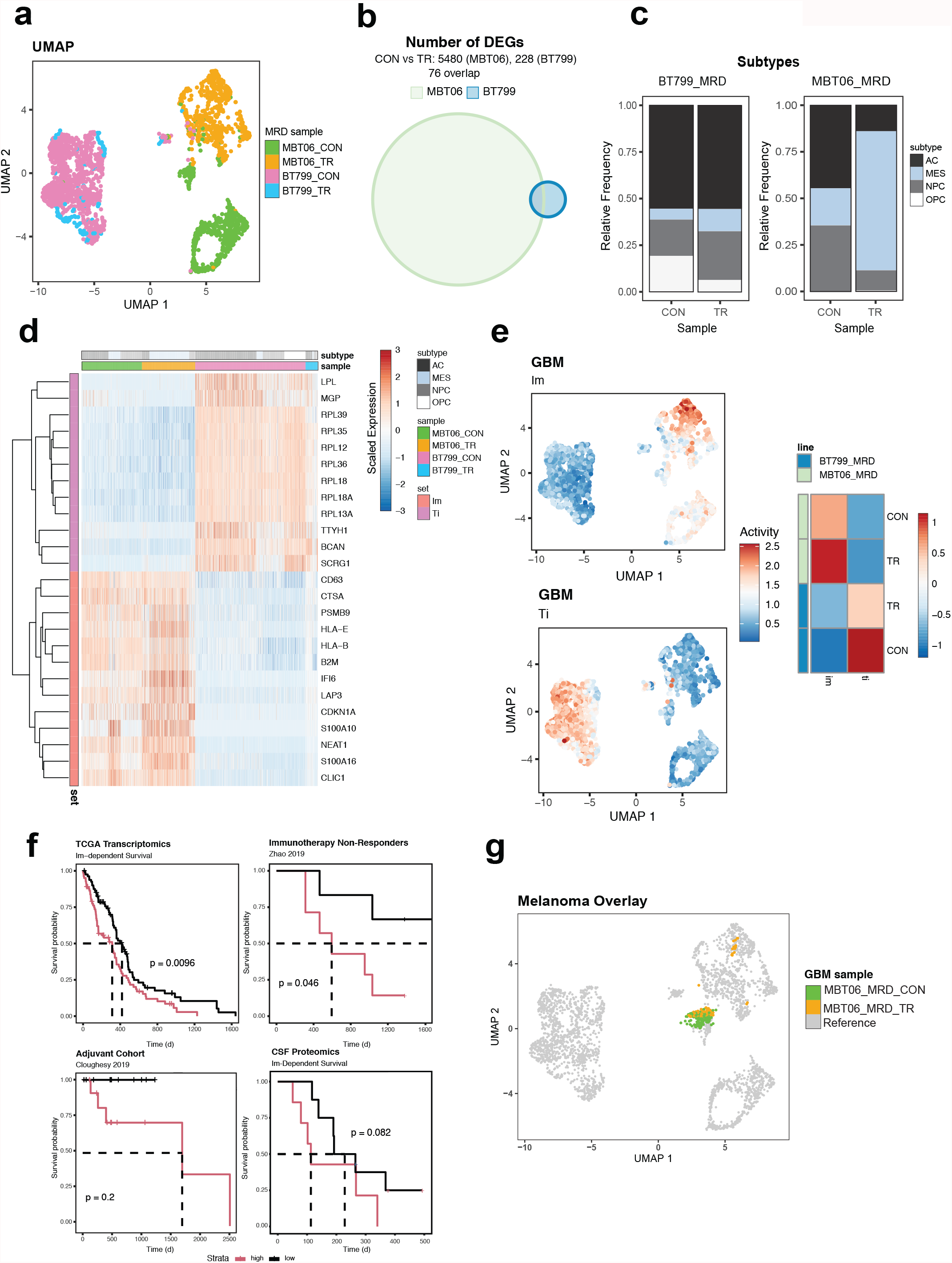
scRNAseq reveals prognostic immunomodulatory signature in GBM patients. **a**. UMAP of MBT06 and BT799 samples at minimal residual disease (MRD) timepoint in response to chemoradiotherapy (TR; along with time-matched controls, CON). **b**. Venn diagram of differentially-expressed genes between control and treated BT799 and MBT06 cells. **c**. Relative abundance of GBM Subtypes. **d**. Expression heatmap of immunomodulatory (Im) and translation-promoting (Ti) gene panels. **e**. UMAP of Im and Ti activities (*top*) and signature activity heatmap stratified by MRD sample (*bottom*) **f**. Kaplan-Meier survival analysis, stratified into Im^high^ and Im^low^ groups, for (top left) TCGA GBM transcriptomic data, (top right) Zhao *et al*.^23^ immunotherapy non-responders transcriptomic data, (bottom left) Cloughesy *et al*.^21^ adjuvant-therapy treated patient transcriptomic data and (bottom right) CSF GBM proteomic data. Dashed lines are median survival times. **g**. Mapping of melanoma samples onto GBM UMAP space using transfer-learning based method. Melanoma samples are highlighted in color; Colors represent which GBM sample the melanoma cells show the highest degree of similarity to.

Uniform Manifold Approximation and Projection (UMAP)-based embedding revealed three major cellular populations (**Fig 4a, Supp Fig 4c**). The first two distinct populations represented control and treated MBT06 samples (*green* and *orange* populations, respectively) and showed pronounced treatment-induced differences in higher dimensional transcriptomic space (**Fig 4a**) and by differential expression analysis (**Fig 4b**), whereas the third population representing control and treated BT799 cells co-clustered, suggesting a less pronounced response to treatment. These profiles were in agreement with our barcode analysis.

To assess intratumoral heterogeneity, we classified cells into mesenchymal (MES)-, astrocyte (AC)-, oligodendrocyte progenitor (OPC) and neuronal progenitor (NPC)-like subtypes^18^. We found that at least three of four GBM-subtypes were represented in each sample; however, the relative abundance of each subtype was more stable in BT799 compared to MBT06 following treatment (**Fig 4b, Supp Fig 4d**). In untreated BT799 samples, AC-like cells were the most abundant, and the proportion of each state was not significantly affected following treatment (p=0.982, Χ ^2^ test). In contrast, the untreated MBT06 sample was predominantly composed of AC- and NPC-like cells, and treatment induced a significant shift towards the MES-like state (p<0.001, Χ ^2^ test). We also scored control and treated cells using pan-cancer cell-state gene sets^19^ (**Supp Fig 4e**). Notably, treated MBT06 samples were associated with upregulation of quiescence, apoptosis, inflammation and hypoxia signatures, and downregulation of cell cycle and proliferation signatures. By comparison, treatment of BT799 was associated with negligible changes in cancer state activities, corroborating the lack of treatment-induced transcriptomic changes observed in BT799 samples at the global UMAP level.

To identify innate tumor-specific differences at MRD, we performed differential expression analysis of control MBT06 and BT799 cells. MBT06 was enriched for tumor-specific immune signalling including antigen presentation, whereas BT799 was enriched for translation initiation and rRNA processing. To compare the regulation of these pathways and evaluate their clinical utility, we developed an immunomodulatory (Im) and translation-initiating (Ti) signature composed of 13 and 12 genes, respectively, that were differentially expressed between the two samples and coherent across independent GBM datasets^18, 20^, including scRNAseq profiles obtained in the current study (**Fig 4d, Supp Fig5a, b)**. The Im signature consisted of genes comprising the MHC class I protein complex, while the Ti signature was primarily composed of genes expressing large ribosomal subunit (**Supp Fig 5c**). Furthermore, the Im signature was correlated with the expression of mesenchymal-like and injury-response signatures whereas the Ti signature is correlated with the expression of developmental-like signatures (**Supp Fig 5d**). While these signatures were derived from untreated samples, treated MBT06 and BT799 samples consistently responded with an increased expression of the immunomodulatory signature and suppression of the translation-promoting signature (**Fig 4e**).

### Immunomodulatory signature predicts survival in GBM

We next sought to determine whether our immunomodulatory and translation-promoting signatures had prognostic value, as they were derived from the differential pathway activities of tumors associated with the best and worst disease outcomes in our cohort. We evaluated the prognostic value of our signatures using 4 independent GBM transcriptomic datasets [MRD mouse models (N = 5; *current study*), public TCGA-GBM data (N = 162 patients), immunotherapy-responder (N = 13) and non-responder (N = 12) GBM patient data^23^, and adjuvant (N = 15) or neo-adjuvant (N = 13) therapy-treated GBM patient data^21^ and 1 GBM proteomic dataset obtained by profiling the cerebrospinal fluid (CSF) obtained from GBM patients (N = 24; *current study*). Intriguingly, despite the Im signature being derived from a sample (MBT06) associated with the best survival outcome (**Supp Fig 5e**), the Im^High^ status were consistently associated with worse prognoses in TCGA samples (**Fig 4f [top left]**, p = 0.0096; **Supp Fig 6**), immunotherapy non-responders (**Fig 4f [top right]**, p = 0.046), adjuvant-treated patients (**Fig 4f [bottom left]**, p = 0.2) and CSF samples (**Fig 4f [bottom right**], p = 0.082). Importantly, among patients that responded to immunotherapy or those that received neo-adjuvant therapy, the Im signature offered to further prognostic value (p = 0.56 and 0.50, respectively; **Supp Fig 7**). In those datasets we also found that the Im signature was expressed at similar levels in immunotherapy responders and non-responders (**Supp Fig 7b**), and in those treated with adjuvant and neoadjuvant therapy (**Supp Fig 7e**). Unlike the immunomodulatory signature, the translation-initiating signature was not associated with survival in any of the validation sets (**Supp Fig 5e, Supp Fig 6a, b**). The incongruity between Im signature being derived from best surviving MBT06 sample and prognosticating poor-survival in larger GBM cohorts may be due to the presence and absence of intact immune system in GBM patients and PDX models, respectively, resulting in differential interaction between Im transcriptomic program and survival.

To further validate the prognostic value of our signatures, we applied our signatures to a comparable single-cell RNAseq dataset from post-treatment, patient-derived melanoma xenografts profiled at MRD^14^. Although clinical data was lacking, we found that MRD melanoma xenograft models that were matched to our control and treated MRD GBM samples (see methods for details) exhibited comparable increases and decreases in immunomodulatory and translation-initiation signatures, respectively, which was consistent with the treatment-response patterns identified in our own GBM samples (**Supp Fig 5f**). Finally, despite GBM being widely recognized as an immunologically “cold” tumor, we found that 45% of cells in melanoma, which are considered immunologically “hot” tumors, mapped onto the control MBT06 transcriptomic space, suggesting that MBT06 samples exhibit characteristics of an immunologically “hot” tumor, and further validates our immunomodulatory signature which was derived from MBT06 and not BT799 (**Fig 4g**).

Taken together, Im-dependent signalling offers a reliable biomarker in GBM across all stages of disease at the bulk and single-cell transcriptomic, and CSF proteomic level, and putatively in other solid cancers.

## Discussion

In this study, we used a high-complexity barcoding library to assess differential patterns of recurrence in response to SoC therapy using five patient-derived GBM xenograft models. Our findings suggest that treatment imposes a selective pressure that enriches for clones that eventually dominate recurrent tumors. Single clone tracking from engraftment through to recurrence provided insight into intratumoral variations of clonal expansion, particularly from MRD to recurrence. Previous study on GBM clonal dynamics in response to chemotherapy alone showed that pre-existing chemo-resistant clones facilitate GBM recurrence^24^. Since then, studies have demonstrated that treatment itself can induce changes in cell populations that then give rise to treatment-resistant tumors^11,25^. Our findings expands on this work and suggest that patterns of GBM recurrence following combined chemoradiotherapy exist on a spectrum between *a priori* fitness, whereby distinct pre-existing clones remain abundant from engraftment through to recurrence, and *a priori* equipotency of clones, whereby treatment-derived subclonal events dominate clonal composition at each timepoint.

To investigate inter-tumoral heterogeneity observed in clonal dynamics from engraftment to recurrence, we used our therapy-adapted, patient-derived xenograft model to capture MRD, which after pruning of cellular subpopulations by therapy, emerged as the least heterogeneous state with the lowest possible clonal diversity across different patient samples. Single-cell RNA sequencing of MRD samples from the longest and shortest surviving patients in our sample cohort revealed profound cellular ITH in signalling and cell states, independent of treatment. We observed regulation of distinct molecular pathways in each sample, and we derived a prognostic immunomodulatory (Im) signature that was validated across five independent datasets (four transcriptomic and one proteomic). Importantly, the Im signature was found to have significant prognostic value when applied to the TCGA GBM dataset, and datasets published by Zhao *et al*.^23^ and Cloughesy *et al*.^21^ as Im^High^ transcriptomes were associated with worse overall survival. While promising, the requirement for surgically-resected tissues and transcriptomic profiling might limit the clinical utility of our Im signature. Therefore, we assessed the prognostic value of the Im signature in CSF proteomes from GBM patients, and in this independent cohort of 24 patients found the signature predicted worse survival. Finally, we compared our thirteen-gene signature to recently published work by Richards *et al*.^20^ While we found similarity with their >4000 gene injury response signature characterized by inflammation and immune cell activation, we suggest the thirteen-gene signature would be more accessible for clinical application.

It was intriguing to find that Im^High^ GBM cells (MBT06 samples) closely resemble melanoma cells profiled at the MRD stage (**Fig. 4i**). This finding suggests that tumors with high immunoregulatory signalling may resemble immunologically “hot” tumors, like melanoma, and may represent a subset of GBMs that have developed mechanisms to overcome their immunosuppressive niche^22-23^. Taken together, Im-dependent signalling is a useful biomarker in GBM across all stages of disease at the bulk, single-cell transcriptomic and proteomic level, and putatively in other solid cancers.

Profiling the state of MRD has added immeasurable value to the predictive and personalized management of liquid cancers such as Acute Myeloid Leukemia (AML), where sequential molecular MRD monitoring has been used to anticipate relapse, refine clinical decision making and personalize treatment plans^13^. Whole-genome sequencing and mutational integration of plasma cell-free DNA allow ultra-sensitive detection of mutations in MRD to be achieved even in patients with low-burden solid cancers to guide clinical interventions^24^. Characterization of MRD through liquid biopsy, and in the case of brain tumors through interrogation of the CSF^27^, may help to define which solid cancer patients are at risk of relapse after surgery and warrant prophylactic adjuvant treatment^28^, and clinical trials are underway to determine if the predictive value of MRD for recurrent disease guides therapy choices that reduce the risk of recurrence of solid cancers^29, 30^. Excitingly, patients enrolled in certain clinical trials for recurrent GBM are now given intraventricular reservoirs to bypass the blood-brain barrier and allow for locoregional delivery of new immunotherapies^31, 32^, which could present exceptional opportunities to survey the CSF of patients after completion of SoC therapy, at the presumed point of MRD, and interrogate prognostic markers, like the Im signature reported here, to inform post-SoC management. For example, patients with Im^High^ signatures could be stratified to ongoing immunotherapy trials (NCT04201873).

Advantages to treating patients at MRD rather than waiting for clinical relapse to initiate further therapy include the fact that a cancer characterized by vast ITH will be at its most homogeneous state and potentially targetable by a tailored drug regimen against a rare minority of cancer stem-like cells that are now enriched at MRD. In addition, patients are generally clinically well at MRD, and more capable of tolerating drugs with significant side effects compared to a time when they are weakened by fulminant relapse.

Our work applies barcoding and single-cell sequencing technologies toward the first functional, evolutionary and cell state characterization of MRD in GBM, unlocking a new means of predictive and personalized profiling that has, until now, been impossible to interrogate in the course of natural history and disease progression of GBM in patients. Application of our findings through CSF surveillance in clinical trials going forward could aim to implement iterative detection, profiling and targeting of MRD in patients with recurrence of solid cancers, which could anticipate and even lead to prevention of solid cancer recurrence.

## Figure Legends

**Supplementary Figure 1. *In vitro* stem cells functional assays of GBM cell lines in response to chemoradiotherapy**.

a. Sphere forming frequency of five patient derived GBM samples (N=5, mean ± SEM).

b. Limiting dilution assay to determine stem cell frequency in five patient derived GBM samples. Each point represents a technical replicate (N=3).

c. Representative flow plot for gating strategy on the sorting of GFP+ cells from GBM cell lines transduced with barcode library.

**Supplementary Figure 2**.

a-c. Shannon Index (left) displays relative diversity of barcodes in three patient derived GBM cell lines, BT935 (a), BT954 (b), and MBT06(c), through timepoints of *in vivo* modeling and clonal frequency (middle) at the time of recurrence and (right) at MRD in response to chemoradiotherapy. Each curve represents an independent experimental replicate (N=3).

d. Visual representation of the clonal frequency of barcodes in BT428 at the time of recurrence endpoint to be retransplanted and subjected to an additional regimen of chemoradiotherapy.

e. Visual representation of the clonal frequency of barcodes in BT799 at the time of control endpoint retransplanted and subjected to an additional regimen of chemoradiotherapy.

**Supplementary Figure 3**.

a-b. Bubble plot displaying frequency of barcodes in BT935 at the recurrence (a) and MRD (b) timepoints. Each vertical line represents an independent experimental replicate for the timepoint (N=3)

c-d. Bubble plot displaying frequency of barcodes in BT954 at the recurrence (c) and MRD (d) timepoints. Each vertical line represents an independent experimental replicate for the timepoint (N=3)

e-f. Bubble plot displaying frequency of barcodes in MBT06 at the recurrence (e) and MRD (f) timepoints. Each vertical line represents an independent experimental replicate for the timepoint (N=3)

g. Barcode correlation of timepoints throughout in vivo chemoradiotherapy regimens in BT935.

h. Barcode correlation of timepoints throughout in vivo chemoradiotherapy regimens in BT954.

i. Barcode correlation of timepoints throughout in vivo chemoradiotherapy regimens in MBT06.

j. Correlation of barcode abundance among different timepoints of chemoradiotherapy in BT428.

k. Correlation of barcode abundance among different timepoints of chemoradiotherapy in BT799.

**Supplementary Figure 4**.

**a**. UMAP representation of scRNAseq GBM at MRD using lenient filters (*left*; gene/cell > 200, mitochondrial content < 60%) and stringent filters (middle; gene/cell > 3000, mitochondrial content < 10%). *Right*: Cells that passed stringent filter are highlighted in UMAP derived from cells that passed lenient filter.

**b**. Comparison of pseudobulk scRNAseq and bulk RNAseq expression profiles across all profiled GBM samples. *Dashed line*: line of equality; *solid line*: loess curve. UMAP representation of scRNAseq GBM at MRD data, stratified by cluster.

**c**. UMAP representation of GBM at MRD cell clusters.

**d**. BT799 and MBT06 GBM subtype representation plot, colored by treatment status.

**e**. Heatmap of CancerSEA cell state activities in MBT06 and BT799 samples at MRD.

**Supplementary Figure 5**.

**a-b**. Correlation of genes with signature scores at each iteration of Im signature derivation. 31 genes were nominated for Im signature based on FDR = 5% and logFC > 1.5 differential expression between control MBT06 and BT799 samples and were subsequently pruned to 14 genes following iteration 1 (a) and 13 genes following iteration 2 (b). *Left, middle* and *right* plots represent correlation plots for scRNAseq data from current study, Neftel *et al*.^18^, and Richards *et al*.^20^, respectively. Red dashed line: coherence threshold = 0.1; black dashed line: Pearson correlation = 0.

**c**. Hypergeometric overrepresentation analysis of Im and Ti gene signatures using GO biological process (*top*), cellular compartment (*middle*) and molecular function (*bottom*) gene panels.

**d**. Correlation heatmap showing similarity between Im and Ti signatures and GBM-related signatures from Neftel *et al*.^18^ (AC; astrocyte-like; MES; mesenchymal-like, OPC; oligodendrocyte-progenitor-like, NPC; neuroprogenitor-like), Richards *et al*.^20^ and Verhaak *et al*. 2010 using expression profiles from current study (*left*), Neftel *et al*.^18^ 2019 (*middle*) and Richards *et al*.^20^ (*right*).

**e**. Correlation between bulk RNA-estimated Im (*left*) and Ti (*right*) GSVA scores and survival for five GBM PDX models.

**f**. Progression of Im (*left*) and Ti (*right*) gene signatures between control and treated GBM and melanoma samples.

**Supplementary Figure 6**.

**a-b**. Kaplan-Meier survival analysis, stratified into Ti^high^ and Ti^low^ groups, for (a) TCGA GBM transcriptomic data, and (b) CSF GBM proteomic data. Dashed lines are median survival times.

**c**. Survival sensitivity analysis for Im signature using TCGA dataset. For each gene comprising Im signature, Kaplan-Meier survival analysis was performed stratifying GBM samples into gene^High^ and gene^Low^ groups to evaluate the prognostic value of each individual gene, and alternatively Im^High^ and Im^Low^ statuses were computed in the absence of the gene to determine the influence of the gene on the overall prognostic value of the signature.

**Supplementary Figure 7**.

**a**. Venn diagram evaluating overlap between differentially-expressed genes in Responder vs. Non-responder transcriptomic profiles (Zhao *et al*.^23^) and Im signature.

**b**. Relative proportion of immunotherapy responders and non-responders stratified by Im status (high vs. low).

**c**. Kaplan-Meier survival analysis of immunotherapy responders (Zhao *et al*.^23^) stratified into Ag^high^ and Ag^low^ groups. Dashed lines represent median survival times.

**d**. Im signature expression stratified by therapy type (adjuvant vs. neoadjuvant; Cloughesy *et al*.^21^).

**e**. Kaplan-Meier survival analysis of neoadjuvant-treated patients (Cloughesy *et al*.^21^) stratified into Ag^high^ and Ag^low^ groups. Dashed lines represent median survival times.

