## Supplementary material for "Characterization of the minimal residual disease state reveals distinct evolutionary trajectories of human glioblastoma": Supplemental Material - Jan 12.pdf

#### Supplementary Figure 1

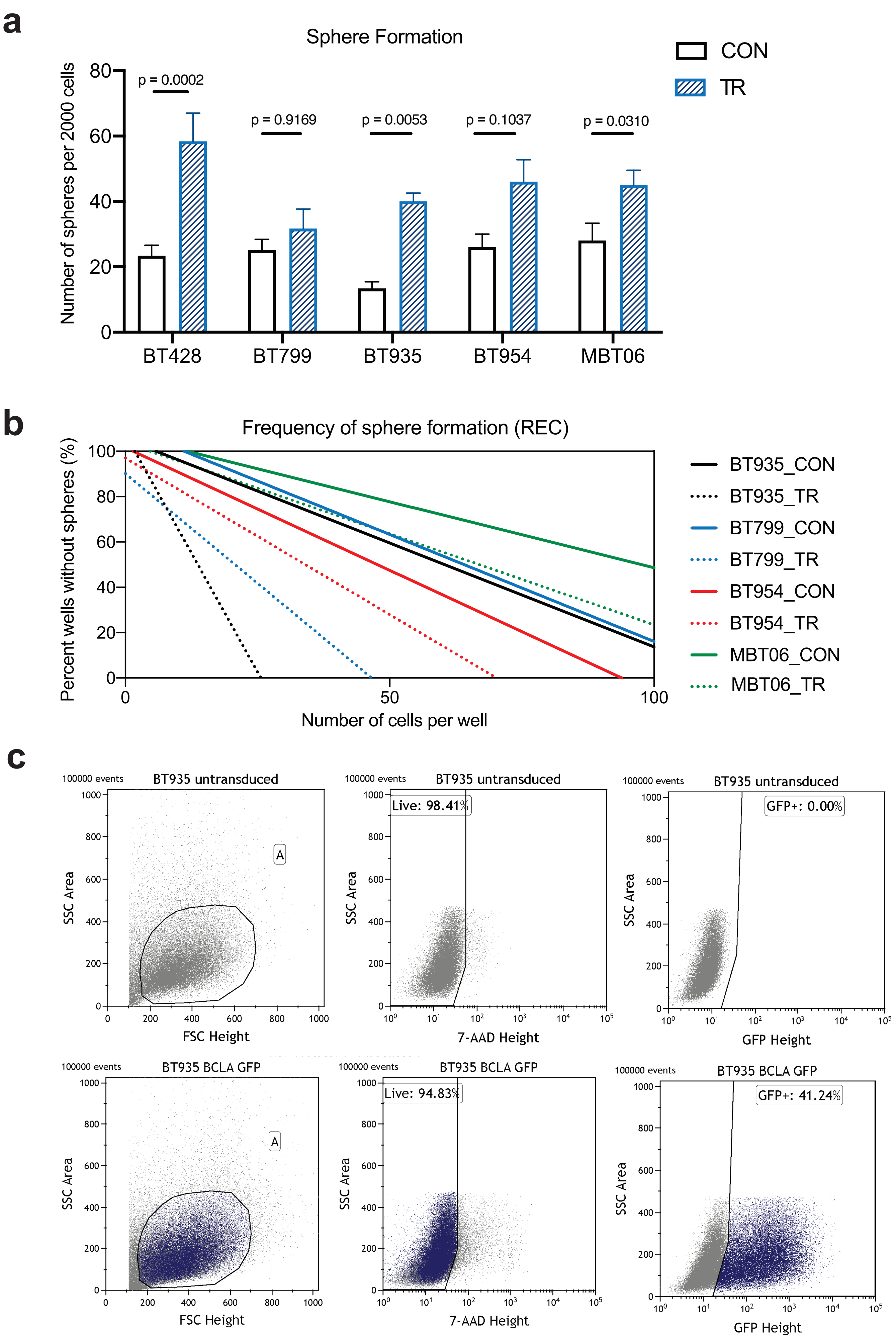

### Supplementary Figure 2

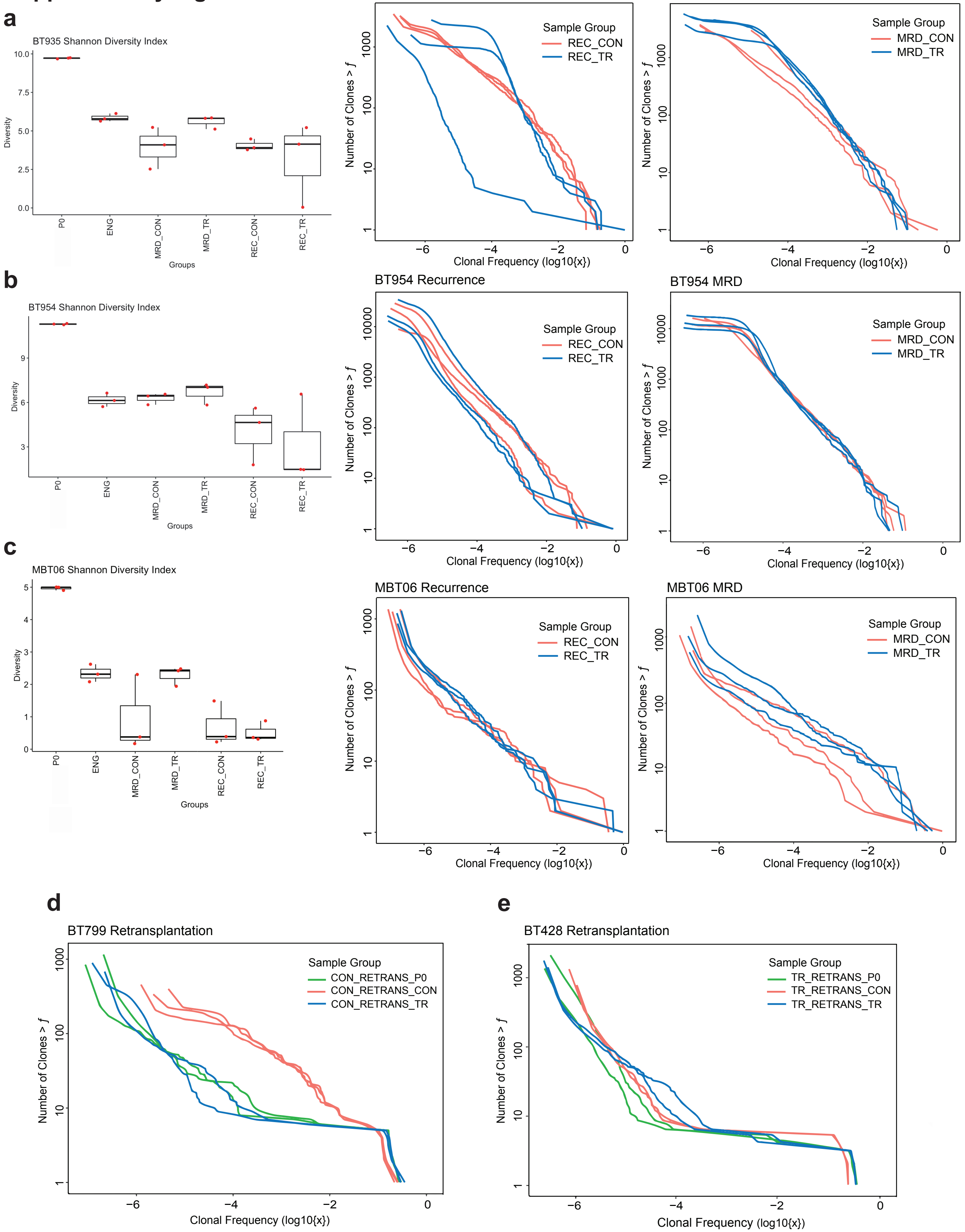

#### Supplementary Figure 3-1

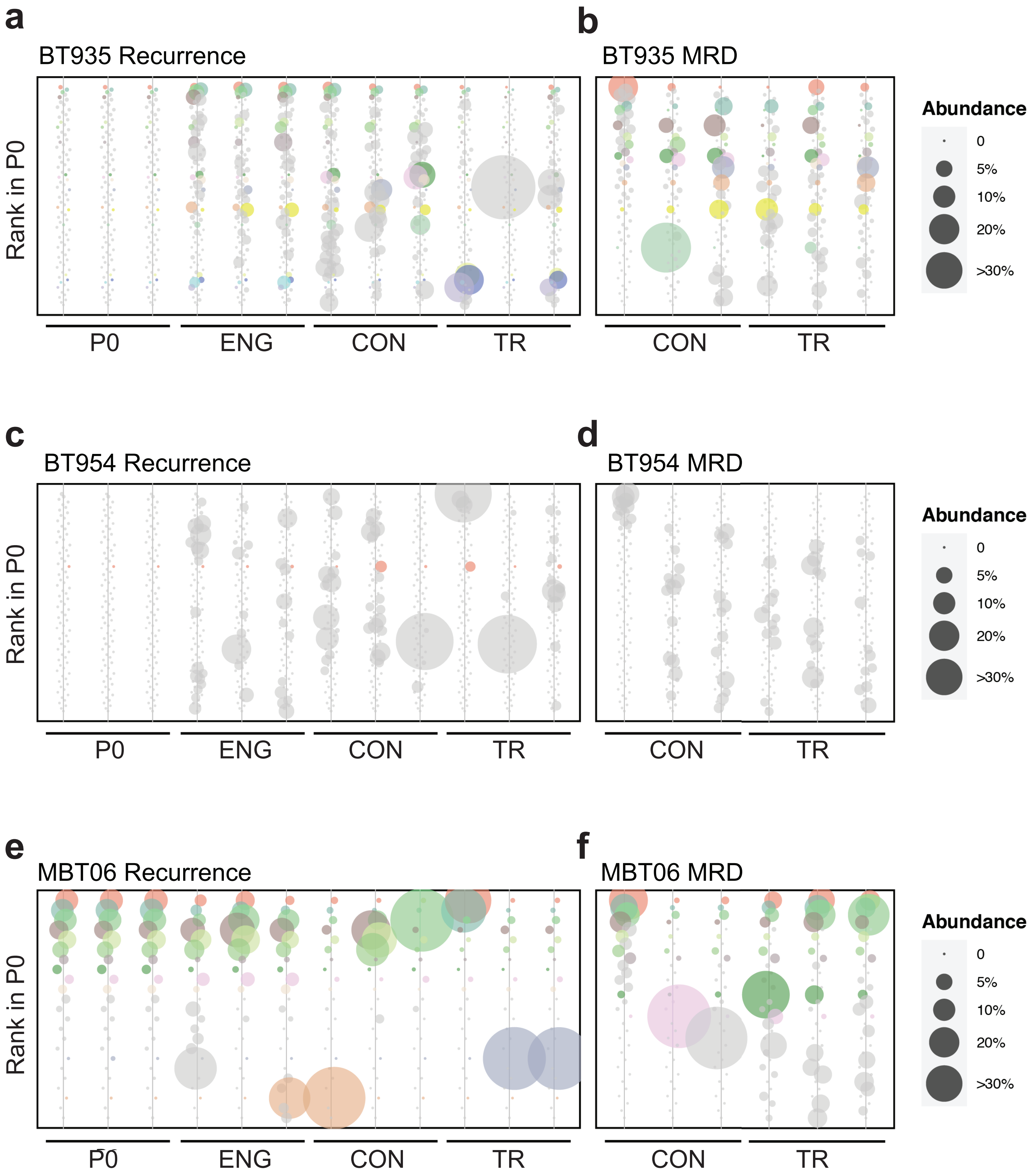

#### Supplementary Figure 3-2

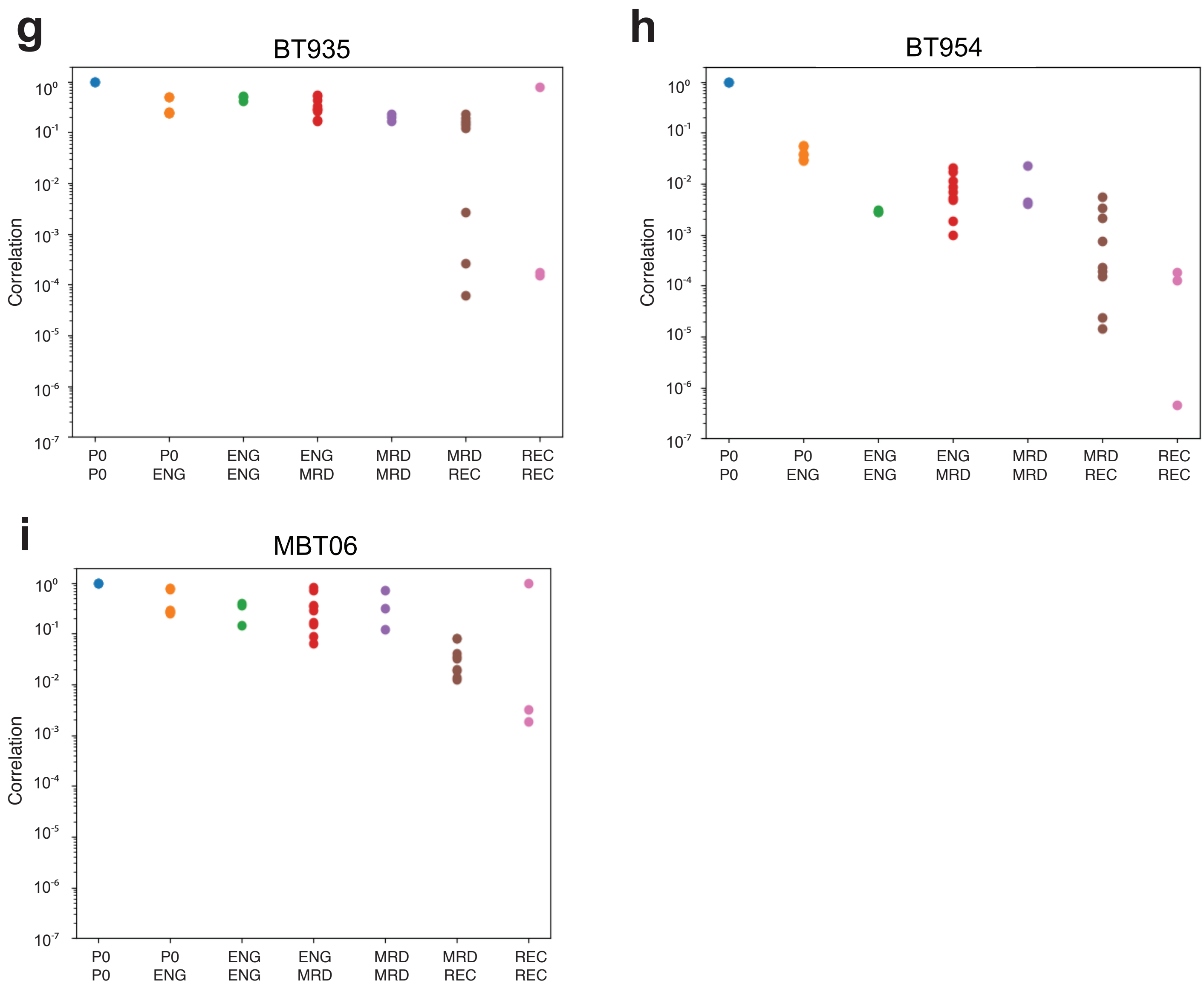

### Supplementary Figure 3-3

**j**

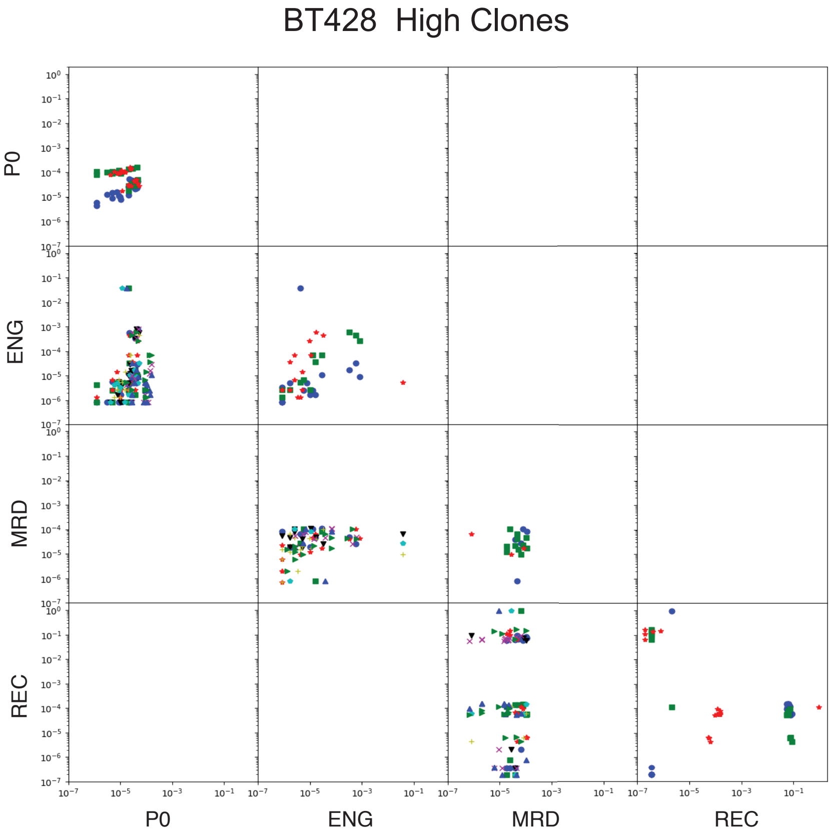


**k**

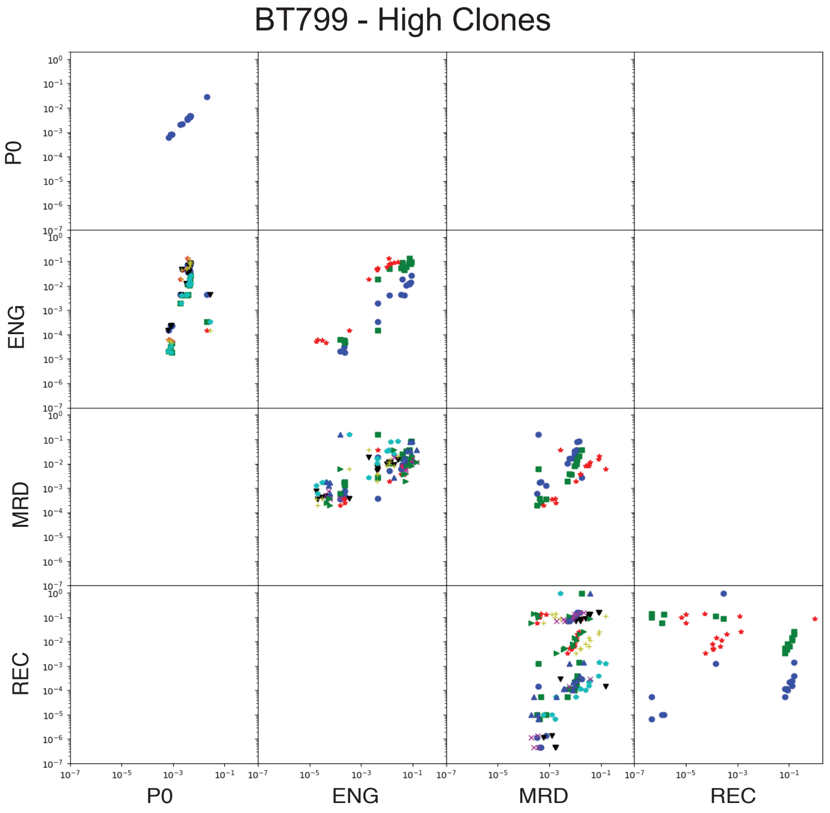

### Supplementary Figure 4

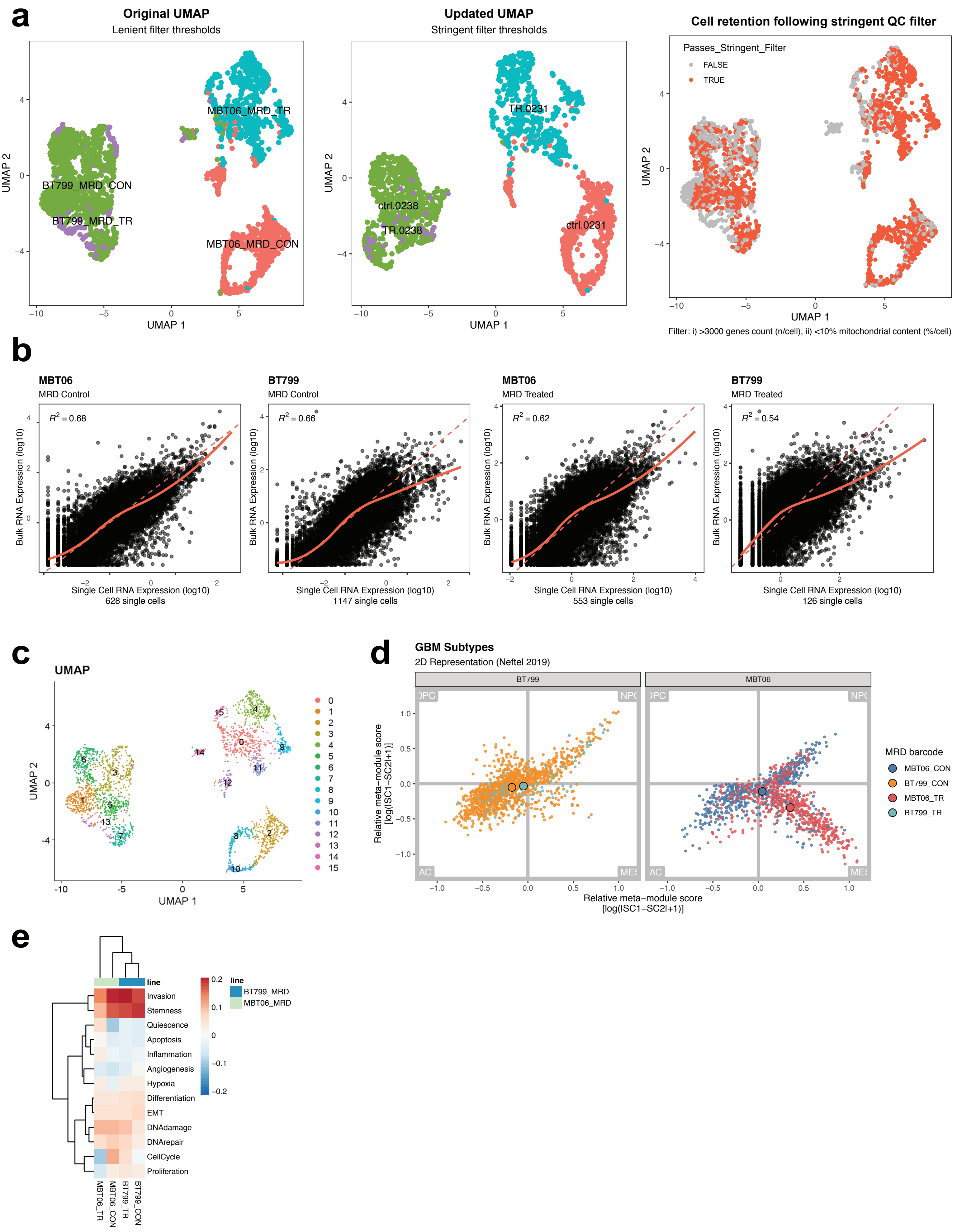

### Supplementary Figure 5

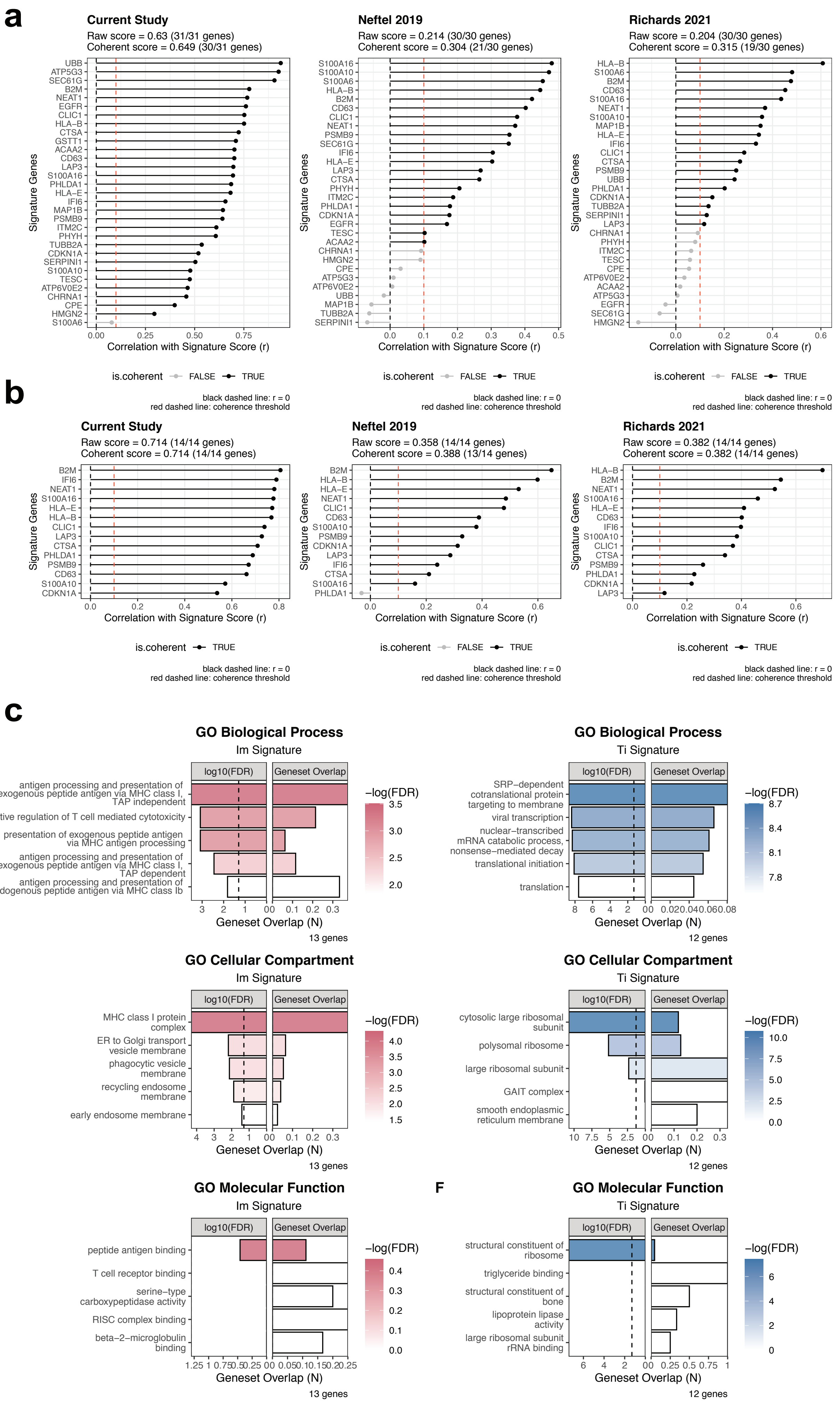

### Supplementary Figure 5-2

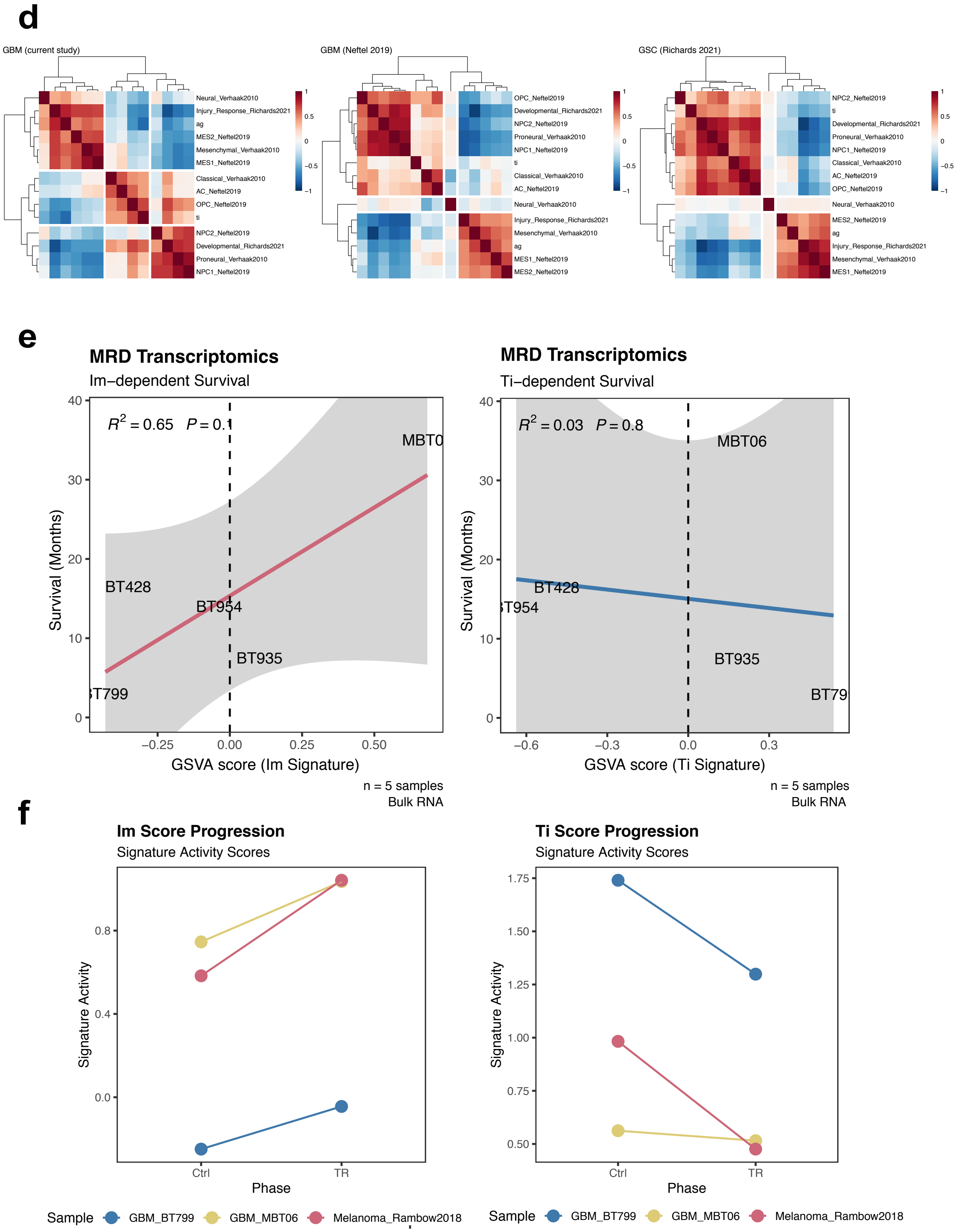

### Supplementary Figure 6

**a**

**b**

**c**

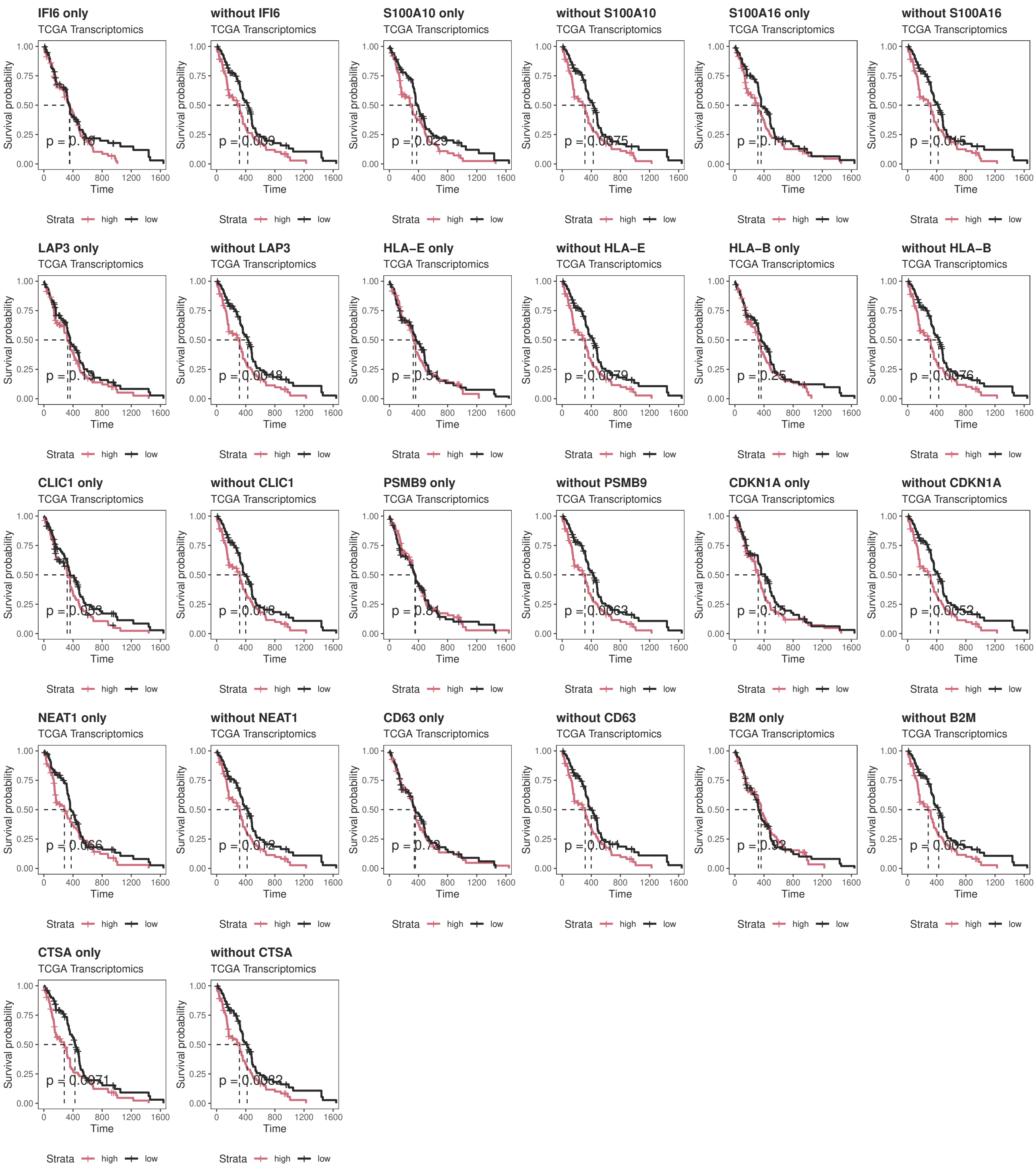

### Supplementary Figure 7

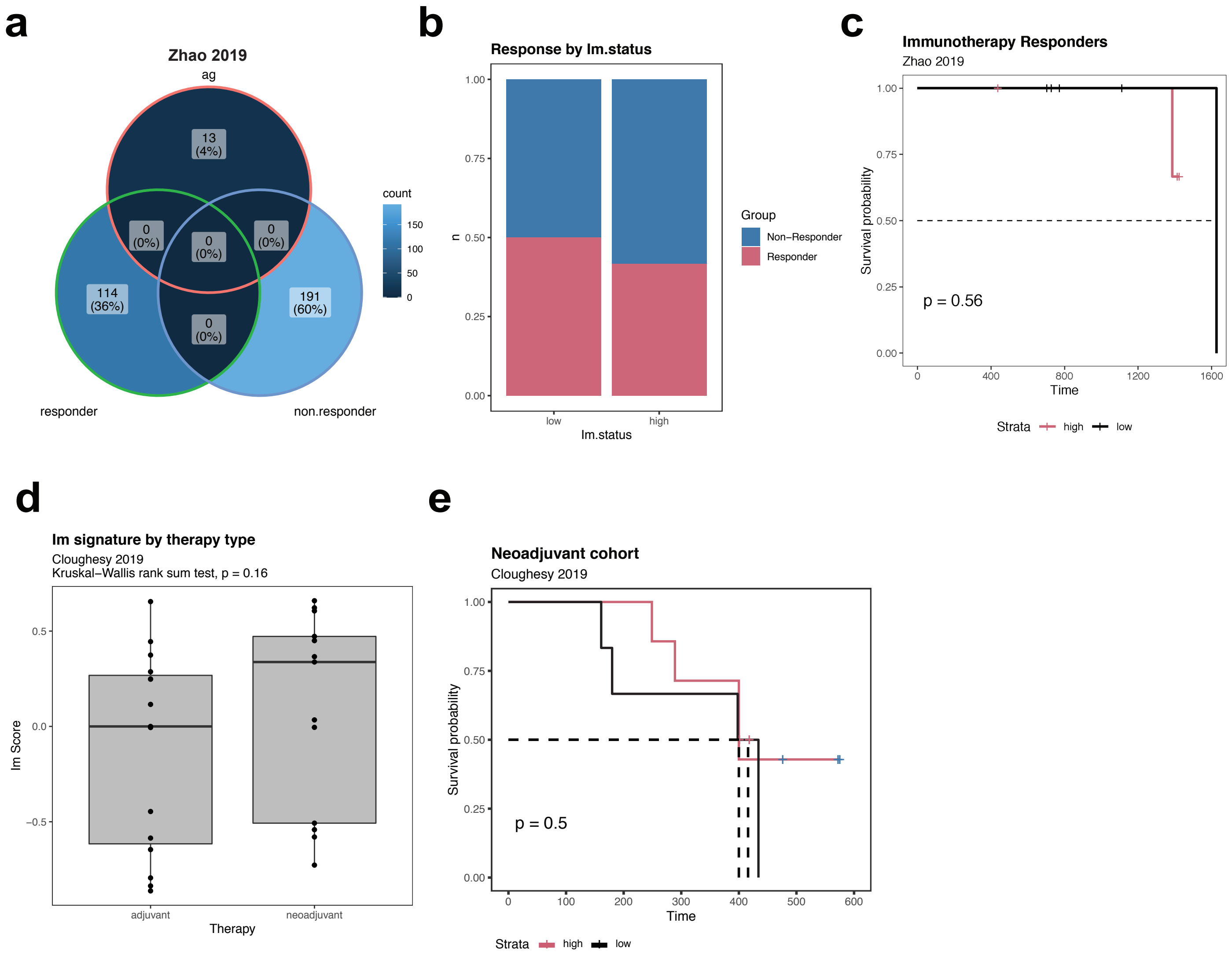

#### Supplementary Table 1

| Sample | Age/Sex | Diagnosis | Subtype (Bulk RNA) | Survival (Months) | MGMT Expression | TP53 Expression | P53 Variants | CDKN2A | RB1 Expression | IDH Status |
|---|---|---|---|---|---|---|---|---|---|---|
| *BT428* | 63/F | GBM | Proneural | 14 | - | + | | - | + | Wildtype |
| *BT799* | 77/F | GBM | Proneural | 3 | - | + | V157D | +/- | + | Wildtype |
| *BT935* | 53/F | GBM | Mesenchymal | 8 | - | + | E286K | +/+ | + | Wildtype |
| *BT954* | 65/F | GBM | Classical | 15 | + | + | | - | - | Wildtype |
| *MBT06* | 50/F | GBM | Mesenchymal | 43+ | - | + | | - | + | Wildtype |

#### Supplementary Table 2

| Treatment | P0 | Engraftment (ENG) | Minimal Residual Disease (MRD) | Recurrence (REC) |
|---|---|---|---|---|
| *Control (CON)* | BT428<br>BT799<br>BT935<br>BT954<br>MBT06 | BT428<br>BT799<br>BT935<br>BT954<br>MBT06 | BT428<br>BT799<br>BT935<br>BT954<br>MBT06 | BT428<br>BT799<br>BT935<br>BT954<br>MBT06 |
| *Temozolomide (TMZ)* | | | BT428<br>BT799<br>BT935<br>BT954<br>MBT06 | BT428<br>BT799<br>BT935<br>BT954<br>MBT06 |
| *Radiation Only (R)* | | | BT428<br>BT799<br>BT935<br>BT954<br>MBT06 | BT428<br>BT799<br>BT935<br>BT954<br>MBT06 |
| *Temozolomide and Radiation (TR)* | | | BT428<br>BT799<br>BT935<br>BT954<br>MBT06 | BT428<br>BT799<br>BT935<br>BT954<br>MBT06 |

Bulk RNA Sequencing and barcoding
Bulk RNA Sequencing, scRNAseq and barcoding

#### Supplementary Table 3

| Sample | CON | TR | P-value |
|---|---|---|---|
| *BT428* | 174 | 295 | 0.0246 |
| *BT799* | 59 | 160 | 0.0177 |
| *BT935* | 90 | 247 | 0.0246 |
| *BT954* | 194 | 199 | 0.832 |
| *MBT06* | 160 | 215 | 0.1098 |
