## Supplementary material for "Characterization of the minimal residual disease state reveals distinct evolutionary trajectories of human glioblastoma": Materials and Methods

### *Dissociation and culture of primary GBM tissue*

Human GBM samples (**Supp. Table 1**) were obtained from consenting patients, as approved by the Hamilton Health Sciences/McMaster Health Sciences Research Ethics Board. Brain tumor samples were dissociated in PBS containing 0.2 Wünsch unit/mL Liberase Blendzyme 3 (Roche), and incubated in a shaker at 37˚C for 15 min. The dissociated tissue was filtered through a 70µm cell strainer and collected by centrifugation (1500 rpm, 3 min). Red blood cells were lysed using ammonium chloride solution (STEMCELL Technologies). GBM cells were resuspended in Neurocult complete (NCC) media, a chemically-defined serum-free neural stem cell medium (STEMCELL Technologies), supplemented with human recombinant epidermal growth factor (20ng/mL: STEMCELL Technologies), basic fibroblast growth factor (20ng/mL; STEMCELL Technologies), heparin (2 µg/mL 0.2% Heparin Sodium Salt in PBS; STEMCELL technologies), antibiotic-antimycotic (1X; Wisent), plated on ultra-low attachment plates (Corning) and cultured as neurospheres.

### *Propagation of brain tumor stem cells (BTSCs)*

Neurospheres derived from minimally-cultured human GBM samples were plated on polyornithine- laminin coated plates for adherent growth. Adherent cells were replated in low-binding plates and cultured as tumorspheres, which were maintained as spheres upon serial passaging *in vitro*. These cells retained their self-renewal potential and were capable of *in vivo* tumor formation.

### *Self-renewal neurosphere formation assay*

Once primary neurosphere formation was noted in GBM cultures, neurospheres were dissociated to single cells and re-plated in 0.2 mL NSC media as previously published[1]. Briefly, neurospheres were treated with Liberase and plated at 200 cells/well density in a 96-well microwell plate in 0.2 mL volume of NSC media. The spheres were counted seven days later.

### *In vivo therapy model*

$1x10^6$ GBM cells (BT428, BT799, BT935, BT954 and MBT06) were injected in NOD.Cg-*Prkdc*$^{scid}$ *Il2rg*$^{tm1Wjl}$/SzJ (NSG) mouse brains as previously described[1]. Briefly, mice were anesthetized using gas anesthesia (Isoflurane: 2.5%). Using a 15-blade scalpel a 1.5 cm vertical midline incision was made on top of the skull. A small burr hole was then made (2-3 mm anterior to the coronal suture, 3 mm lateral to midline) using a drill held perpendicular to the skull. A Hamilton syringe was used to inject 10µL of cell suspension of GBM cells into the frontal lobe. The syringe was inserted through the burr hole at a 30$^{o}$ angle to a 5 mm depth. The incision was closed using interrupted stitches and sutures were sealed with a tissue adhesive. Mice were identified using ear notches and placed in recovery cages. Mice were monitored weekly for signs of illness and sacrificed at endpoint.

Once tumor engraftment was confirmed using MRI, mice were randomly assigned to control or treatment groups and chemoradiotherapy treatment was started as described in Fig 1a. For mice in the combined chemoradiotherapy arm, 50 mg/kg TMZ was oral gavaged one hour before cranial irradiation was administered to mice. The mouse shield is designed such that only the head of the mice is irradiated.

### *Lentiviral production and GBM transduction of barcode library (BCLA)*

Oligonucleotides comprising a 12 base pair degenerate region (the barcode) followed by two stable bases (C or G), and one of several four base pair library codes, were synthesized with common flanking regions (Sigma Aldrich, St. Louis, MO, USA). Nested PCR using the common regions generated double-stranded DNA, which was subsequently ligated into a second-generation lentiviral vector (pLJM1) containing puromycin resistance cassette and ZsGreen fluorescent marker. The barcode library construct was packaged into lentiviruses as follows: BCLA plasmid (6 µg), psPAX2 (6.0 µg) and pMD2.G (4.0 µg) plasmid was transfected into HEK293FT cells using lipofectamine 2000. The viral suspension obtained after 48 hours was precipitated using PEG-it (System Biosciences) and the pellet was suspended in NSC media, aliquoted and frozen at -80°C.

The viral titer of BCLA was determined individually for all five primary GBM cell lines. Briefly, 140,000 cells were plated in 200µL of NSC media with multiple dilutions of BCLA lentivirus (1-, 200-, 2000 and 20,000-fold dilutions). After 24 hours, the media was changed, and cells were left for an additional 24 hours. The percentage of GFP+ cells in each dilution for each cell line was determined using flow cytometry.

Treatment-naïve GBM cell lines were transduced using the BCLA library at a low multiplicity of infection (MOI $\leq$ 0.3), such that, on average each cell was uniquely barcoded. Barcoded cells were sorted by GFP expression (**Supp Fig 1c**) and expanded *in vitro* prior to intracranial transplantation in immunodeficient mice.

To assess clonal composition, we collected mouse brains at three time points (n=3 per time point), corresponding to ENG, MRD and REC of chemoradiotherapy-treated mice (TR). Vehicle-treated control (CON) mice were also collected at matched time points at MRD (MRD_CON), and at disease endpoint (REC_CON). The reference clonal composition was also examined prior to intracranial injection (P0). At the experimental endpoints, (i.e. REC_CON and REC_TR), samples from two GBM models (i.e. BT428 and BT799) were serially transplanted, and barcode distributions were examined in the re-engrafted mouse cohort following outgrowth of five to seven days (**Fig 1b**).

### *Genomic DNA extraction and PCR amplification*

Gentra Puregene Tissue kit was used to extract all genomic DNA from full mouse brains. Considering that the GBM tumor population in a mouse brain is expected to be significantly smaller, especially at the MRD timepoint, we optimized genomic DNA extraction protocols for snap-frozen mouse brain samples and performed nested-PCR to amplify the 137 bp barcode region for genomic sequencing. Spike-in control barcoded HEK293 cells were added to mouse brain samples for gDNA extraction to estimate clone size. We used NEB UltraQ II polymerase to amplify the 137 bp barcode regions using nested-PCR, which were then visualized on a 3% agarose gel. The primer sequences are in (**Supp Table 3**).

### *Barcode processing and analysis*

FASTQ files for each sequenced sample were processed using a bespoke Perl script. Each read was examined to identify one of the three expected library codes (CCAA, ACGT, or TGGA) followed by eight bases corresponding to the vector sequence (eg: ATCGATAC), allowing up to one mismatched base for each feature. Reads lacking both of these sequences were discarded. The nucleotide sequence corresponding to the barcode was then extracted as the 18 nucleotides preceding the vector sequence, and all unique barcodes were counted. All barcode count files, one per sample, were then merged into a single matrix. Noise introduced through sequencing or PCR errors was reduced by collapsing barcodes within a Hamming distance of two into a single barcode record, where the barcode with the highest average abundance was retained as the "parent" barcode.

Next, samples with fewer than 100000 filtered sequence reads were discarded, and each sample was normalized for sequencing depth by dividing all read counts in a given sample by the sum of total reads. Technical replicate samples were combined by summing counts across replicates, and barcodes that were observed in only one sample were removed as potential artefacts. The final barcode matrix comprised 1,170,776 barcode sequences.

### *Bioinformatic Analysis*

Barcode analysis and visualization was performed in R (v4.0.0) unless otherwise indicated. The Shannon Diversity Index of the barcode populations was computed using the 'diversity()' function from the 'vegan' R package (v.2.5-6). The center line represents the median, the box limits represent the upper and lower quartiles; whiskers represent the 1.5x interquartile range, and each dot represents an individual biological replicate. Bubbleplots were generated using ggplot2 (v3.3.2) using barcodes that enrich beyond 1% in at least one sample.

### *Correlation analysis*

To understand how individual clones travelled across the different sampled time points, we computed intra-sample and inter-sample Pearson correlation analysis, with each point representing a comparison of independent biological replicates. To further focus on the clones that are large in REC samples, we traced their abundance in each of the earlier timepoints.

### *Single-cell RNA sequencing analysis*

Preprocessing.

Raw FASTQ files were aligned to the hg19 genome using STAR aligner (STAR v2.5.2b), and the CellRanger (v2.1.1) pipeline was used to obtain the filtered gene-barcode count matrix. Raw count matrices were loaded into a Seurat object, and filtered to retain cells with (*i*) 200 – 9000 recovered genes per cell and (*ii*) less than 60% mitochondrial content. To ensure these filters were robust to poor quality cells, we compared results with a subset of the data retaining only cells >3000 genes per cell and <60% mitochondrial content. To normalize expression values, we adopted the modeling framework previously described and implemented in the sctransform R Package[2]. In brief, count data were modelled by regularized negative binomial regression, using sequencing depth as a model covariate to regress out the influence of technical effects, and Pearson residuals were used as the normalized and variance stabilized biological signal for downstream analysis.

Dimension reduction and clustering.

Dimension reduction was performed using principal component analysis, and the 30 top principal components were retained for UMAP embedding (max_components = 2, n_neighbours = 50, min_dist = 01, metric = cosine)[3]. To identify sub-populations, we performed clustering using the Louvain algorithm implemented in the Seurat package (resolution = 1). Cluster membership was visualized using UMAP.

### *Cell state scoring*

GBM subtype scoring

To classify GBM cells into oligodendrocyte progenitor (OPC), astrocyte (AC), mesenchymal (MES) or neuro-progenitor cell (NPC)-like subtypes, we generated a cell-state representation map using the GBM subtype scoring pipeline described by Neftel and colleagues[4] and compared the proportions of each state in control and treated samples using a chi-square test.

Cancer cell state scoring

To characterize the core cancer state activities in GBM samples, we used the cancer single-cell functional state atlas (CancerSEA) generated by Yuan and colleagues[5]. For each cancer-state gene panel, cells were scored using the AddModuleScore() function in Seurat and scores were averaged across each GBM sample to obtain sample-specific functional state activities.

***Signature Markers***

Differential gene expression

To identify differentially expressed genes across GBM samples, we ran FindMarkers() function in Seurat and applied a FDR < 5% and logFC > 1.5 filter.

Signature derivation

Genes that were differentially-expressed between control MBT06 and BT799 samples were nominated for further gene signature consideration. To derive the gene signature, the expression of each gene comprising the signature had to be positively correlated with the aggregate signature score [computed using the AddModuleScore() function (*Seurat* R package)]. This criterion was enforced across three independent GBM scRNAseq datasets: current study, Neftel et al.[4], and Richards et al.[6]. If the correlation between any given gene and the signature score did not exceed the coherence threshold (r = 0.1), the gene was omitted, and the signature score was recomputed again using the remaining genes. This process was repeated iteratively until convergence upon a stable signature and yielded a MBT06-specific panel (13 genes) and a BT799-specific panel (12 genes). We then performed hypergeometric overrepresentation analysis (*fora* function, *fgsea* R package) using GO biological processes (BP), cellular components (CC) and molecular function (MF) gene sets, and significantly enriched pathways (FDR < 10%) were used to annotate the MBT06- and BT799-specifc gene panels as immunomodulatory (Im) and translation-initiating (Ti) signatures, respectively. Single-cell Im and Ti signature scores were projected onto the UMAP representation or visualized using a heatmap (*pheatmap* R package).

***Signature validation***

Survival analysis

Im and Ti signatures were assessed for external validity using survival outcomes across five independent datasets: 1) MRD bulk RNA-seq data (N = 5 PDX samples); 2) TCGA RNA-seq V2 data from the TCGA PanCancer Atlas was retrieved from the National Cancer Institute (NCI) Genomic Data Commons (GDC) using TCGAbiolinks R package v.2.16.0 (N = 162 primary GBM tumor samples); 3) Transcriptomic data from Zhao et al.[7] that profiled patients that responded (N = 13) or did not respond (N = 12) to immunotherapy; 4) Transcriptomic data from Cloughesy et al.[8] that profiled patients treated with adjuvant (N = 15) or neoadjuvant (N = 13) therapy; 5) CSF proteomics (N = 24 GBM samples). In each dataset Im and Ti signature scores were calculated by GSVA, and survival associations were assessed by linear regression for the MRD samples or by constructing Kaplan-Meier curves (*survminer* R package) for TCGA and CSF samples in which samples were stratified into $Im^{Low}$/ $Ti^{Low}$ (score < 0) and $Im^{High}$/ $Ti^{High}$ (score ≥ 0) groups.

Comparison with public melanoma scRNAseq data

Public scRNAseq data from a study on melanoma at MRD[9] were used to compare the effect of treatment on different solid tumors at MRD. Raw counts for the melanoma dataset were preprocessed using the same pipeline as GBM data. To enable comparison of the melanoma data to GBM, GBM MRD_CON samples were matched to dabrafenib-trametinib (DT)-treated melanoma samples at the timepoint where treated tumors reached an impalpable size (denoted as phase 2), and GBM MRD_TR samples were matched to melanoma samples that continued DT treatment through MRD to the point of recurrence (denoted as phase 3). Im and Ti GSVA scores were computed for each sample and the trends at MRD and following treatment were compared.

Additionally, we adopted the transfer learning approach described by Stuart and colleagues[10] to determine the extent of transcriptomic similarity between melanoma and GBM cells at MRD. Specifically, melanoma cells were mapped onto the GBM UMAP space by projecting the query (melanoma) PCA structure onto the existing reference (GBM) PCA structure. This enabled identification of corresponding cells (i.e., anchors), thereby allowing the mapping of melanoma cells onto the GBM UMAP space and melanoma cell annotation based on GBM cell similarities. Melanoma cells with transfer scores ≥ 0.7 (transfer score range [0,1]) were retained as successfully mapped cells.

### *CSF proteomic collection and processing*

The majority of CSF samples were collected from intracranially-implanted CSF reservoirs. In the case of samples collected from lumbar punctures, an atraumatic 21-gauge spinal needle was used. The volume withdrawn ranged from 5-15mL per collection. Samples were then aliquoted into 1mL polypropylene tubes. Samples were collected for processing within 0-4 hours after collection, during which the samples are stored at room temperature. The tubes were then spun at 2000*g* for 10 minutes at room temperature to remove cellular debris. Supernatants were then maintained in 1mL aliquots and stored at -80°C.

Protein concentrations were determined by BCA assay (Pierce) and a volume equivalent to 25µg of protein was denatured and alkylated with DTT and iodoacetamide, respectively. CSF proteins were purified using an adapted MStern technique[11]. The samples were bound to a PVDF 96-well MStern plate (Millipore) facilitated by a vacuum suction manifold (Millipore). Adsorbed proteins were washed with 100mM ammonium bicarbonate (pH=8) and digested for four hours at 37°C via the addition of 50µL of digestion buffer (5% acetonitrile, 100mM ABC, 1mM $CaCl_2$) containing 2µg of trypsin-LysC protease mixture (Promega). The resultant peptides were eluted from the membranes with 50% acetonitrile, lyophilized and desalted with C18 solid-phase extraction tips prepared in-house. 10µL of purified peptides was spiked with 1µL of indexed retention time (iRT) (Biognosys) peptide standard. Overall, 11µL of peptides were loaded onto a 2cm PepMap Acclaim trap column (Thermo Scientific) using an Easy1000 nanoLC (Thermo Scientific). The peptides were separated and detected along a two-hour reversed-phase gradient using a 50cm EasySpray analytical C18 column coupled by electrospray ionization to a Q-Exactive HF Orbitrap mass spectrometer (Thermo Scientific) operating in a Top 20 data-dependent acquisition mode. The acquired raw data was searched using Maxquant (version 1.6.3.3) against a UniProt complete human protein sequence database (v2020_05) also including yeast invertase (SUC2) and iRT standard peptides. Two missed cleavages were permitted along with the fixed carbamidomethyl modification of cysteines, the variable oxidation of methionine and variable acetylation of the protein N-terminus. Relative label-free protein quantitation was calculated using MS1-level peak integration along with the matching-between-runs feature enabling a 2min retention time matching window. False discovery rate (FDR) was set to 1% for peptide spectral matches and protein identification using a target-decoy strategy. The protein groups file was filtered for proteins detected by a minimum of two peptides and then used to carry out further analysis. Missing LFQ values were imputed with normalized iBAQ intensities[12].


1. (32) Chokshi, C., Savage, N., Venugopal, C. & Singh, S. A Patient-Derived Xenograft Model of Glioblastoma. *STAR Protocols* 1, 100179, doi:10.1016/j.xpro.2020.100179 (2020).
2. Hafemeister, C. & Satija, R. Normalization and variance stabilization of single-cell RNA-seq data using regularized negative binomial regression. *Genome Biol* **20**, 296, doi:10.1186/s13059-019-1874-1 (2019).
3. McInnes, L., & Healy, J. (2018). UMAP: Uniform Manifold Approximation and Projection for Dimension Reduction. ArXiv e-prints
4. Neftel, C. et al. An Integrative Model of Cellular States, Plasticity, and Genetics for Glioblastoma. *Cell* 178, 835-849 e821, doi:10.1016/j.cell.2019.06.024 (2019).
5. Yuan, H. et al. CancerSEA: a cancer single-cell state atlas. *Nucleic Acids Res* 47, D900-D908, doi:10.1093/nar/gky939 (2019).
6. Richards, L.M., Whitley, O.K.N., MacLeod, G. et al. Gradient of Developmental and Injury Response transcriptional states defines functional vulnerabilities underpinning glioblastoma heterogeneity. *Nat Cancer* 2, 157–173 (2021).
7. Zhao, J. et al. Immune and genomic correlates of response to anti-PD-1 immunotherapy in glioblastoma. *Nat Med* 25, 462-469, doi:10.1038/s41591-019-0349-y (2019).
8. Cloughesy, T. F. et al. Neoadjuvant anti-PD-1 immunotherapy promotes a survival benefit with intratumoral and systemic immune responses in recurrent glioblastoma. *Nat Med* 25, 477-486, doi:10.1038/s41591-018-0337-7 (2019).
9. Rambow, F. et al. Toward Minimal Residual Disease-Directed Therapy in Melanoma. Cell 174, 843-855 e819, doi:10.1016/j.cell.2018.06.025 (2018).
10. Stuart, T. *et al.* Comprehensive Integration of Single-Cell Data. *Cell* **177**, 1888-1902 e1821, doi:10.1016/j.cell.2019.05.031 (2019).
11. Berger ST, Ahmed S, Muntel J, Cuevas Polo N, Bachur R, Kentsis A, Steen J, Steen H. MStern Blotting-High Throughput Polyvinylidene Fluoride (PVDF)

Membrane-Based Proteomic Sample Preparation for 96-Well Plates. *Mol Cell Proteomics* **14**, 2814-2823, doi:10.1074/mcp.O115.049650 (2015).

12. Wojtowicz EE, Lechman ER, Hermans KG, Schoof EM, Wienholds E, Isserlin R, van Veelen PA, Broekhuis MJ, Janssen GM, Trotman-Grant A, Dobson SM, Krivdova G, Elzinga J, Kennedy J, Gan OI, Sinha A, Ignatchenko V, Kislinger T, Dethmers-Ausema B, Weersing E, Alemdehy MF, de Looper HW, Bader GD, Ritsema M, Erkeland SJ, Bystrykh LV, Dick JE, de Haan G. Ectopic miR-125a Expression Induces Long-Term Repopulating Stem Cell Capacity in Mouse and Human Hematopoietic Progenitors. *Cell Stem Cell*. 1;19(3):383-96. doi: 10.1016/j.stem.2016.06.008. (2016)
